# Molecular determinants and heterogeneity of tissue-resident memory CD8^+^ T lymphocytes revealed by single-cell RNA sequencing

**DOI:** 10.1101/2020.03.02.973578

**Authors:** Nadia S. Kurd, Zhaoren He, J. Justin Milner, Kyla D. Omilusik, Tiani L. Louis, Matthew S. Tsai, Christella E. Widjaja, Jad N. Kanbar, Jocelyn G. Olvera, Tiffani Tysl, Lauren K. Quezada, Brigid S. Boland, Wendy J. Huang, Cornelis Murre, Ananda W. Goldrath, Gene W. Yeo, John T. Chang

## Abstract

During an immune response to microbial infection, CD8^+^ T cells give rise to distinct classes of cellular progeny that coordinately mediate clearance of the pathogen and provide long-lasting protection against reinfection, including a subset of non-circulating tissue-resident memory (T_RM_) cells that mediate potent protection within non-lymphoid tissues. Here, we utilized single-cell RNA-sequencing to examine the gene expression patterns of individual CD8^+^ T cells in the spleen and small intestine intraepithelial lymphocyte (siIEL) compartment throughout the course of their differentiation in response to viral infection. These analyses revealed previously unknown transcriptional heterogeneity within the siIEL CD8^+^ T cell population at several states of differentiation, representing functionally distinct T_RM_ cell subsets as well as a subset of T_RM_ cell precursors within the tissue early in infection. Taken together, these findings may inform strategies to optimize CD8^+^ T cell responses to protect against microbial infection and cancer.

**One sentence summary:** Here, we applied single-cell RNA-sequencing to elucidate the gene expression patterns of individual CD8^+^ T cells differentiating throughout the course of infection in the spleen and small intestinal epithelium, which revealed previously unidentified molecular determinants of tissue-resident T cell differentiation as well as functional heterogeneity within the tissue-resident CD8^+^ T cell population.

## Introduction

During an immune response to microbial infection, CD8^+^ T cells give rise to distinct classes of cellular progeny with unique migratory and functional properties that coordinately mediate clearance of the pathogen (effector cells) and provide long-lasting protection against reinfection (memory cells). Considerable heterogeneity has been previously described within the long-lived memory T cell pool (*1–3*). While central memory (T_CM_) cells exhibit greater self-renewal and plasticity with the ability to rapidly proliferate and differentiate into secondary effector cells upon reinfection, effector memory (T_EM_) cells provide immediate pathogen control via rapid and potent ‘effector’ function. Moreover, recent studies have revealed additional heterogeneity within the classically defined T_EM_ cell population, including long-lived effector (LLE) cells and peripheral memory (T_PM_) cells, which can be distinguished by distinct surface molecule expression and trafficking properties (*1, 2, 4-6*) (*1-3, 7, 8*). In addition to these circulating memory T cell populations, a non-circulating subset, termed tissue-resident memory (T_RM_) cells, has recently been described (*9*). T_RM_ cells are found in most tissues and positioned at key barrier surfaces, such as the skin and intestinal epithelium, where they play critical roles in limiting early pathogen spread and controlling infection, and also help to control the outgrowth of cancer cells (*10–13*). Whereas heterogeneity within the circulating CD8^+^ T cell memory population has been well characterized, it remains unclear whether the tissue-resident CD8^+^ T cell population might also be comprised of distinct subsets that play unique roles in mediating protective immunity.

Recent studies have begun to illuminate the mechanisms regulating T_RM_ cell differentiation, function, and survival. Activation of naïve CD8^+^ T cells occurs in the spleen or draining lymph nodes, resulting in the upregulation of key transcription factors including Blimp-1 (*14*). Recruitment of activated CD8^+^ T cells to nonlymphoid tissue sites is mediated by chemokine receptors that promote tissue entry, such as CCR9 and CXCR3 (*14–16*). Upon entry to tissue, CD8^+^ T cells undergo transcriptional changes that enforce tissue residency, in part by dampening expression of receptors that promote return to circulation such as CCR7 and S1PR1 (*16*), and begin to direct the T_RM_ cell differentiation program. These changes include upregulation of transcription factors such as Hobit, which, together with Blimp-1, repress genes associated with recirculation, including *Klf2, S1pr1*, and *Ccr7*; and downregulation of the T-box transcription factors T-bet and Eomes, enabling TGF*β* responsiveness (*14–16*). TGF*β* signals within the tissue induce expression of CD103, a key factor for tissue retention, while low levels of T-bet expression are required for IL-15 responsiveness, which plays an important role for long-term survival of T_RM_ cells in some tissues (*17*). However, additional unidentified regulators likely contribute to coordinating the T_RM_ cell differentiation program. Moreover, although it has been shown that T_RM_ cells preferentially arise from KLRG1^lo^ precursors in the circulation, it remains unknown whether all cells that enter the tissue are destined to persist and continue their differentiation into T_RM_ cells, or whether T_RM_ cells are derived from a specific subset of precursors within the tissue at early time points.

Several studies have described a core transcriptional signature that is shared among T_RM_ cells from distinct tissues, highlighting several key transcriptional regulators of T_RM_ cell differentiation, including Hobit, Blimp-1, and Runx3 (*14, 18–21*). However, most of these prior studies have focused on relatively late time points after the T_RM_ cell population has been well established, providing only a snapshot of the gene-expression patterns utilized by T_RM_ cells and potentially missing early events important for their differentiation. Moreover, since these studies relied on RNA sequencing of bulk cell populations, potential heterogeneity representing distinct functional subsets or intermediate states of differentiation may have been missed.

Single-cell RNA sequencing (scRNA-seq) is a powerful approach that can reveal heterogeneity within cell populations and has been used extensively in recent studies to probe the dynamic gene-expression patterns within a wide range of immune cell types in health and disease (*22–31*). This approach has allowed for the elucidation of new cell subsets and states, such as highly efficacious subpopulations of tumor-infiltrating lymphocytes and early states of differentiation for circulating effector and memory CD8^+^ T cells (*22, 24, 25*). Here, we utilized scRNA-seq to generate a single-cell transcriptomic resource dataset elucidating the dynamic gene expression patterns of individual CD8^+^ T cells in the spleen and small intestine intraepithelial lymphocyte (siIEL) compartment throughout the course of their differentiation in response to viral infection. These analyses demonstrate that circulating and tissue-resident memory CD8^+^ T cells utilize highly overlapping patterns of gene expression, revealing a core transcriptional program used by both subtypes of memory CD8^+^ T lymphocytes throughout their differentiation. In addition, these analyses elucidated sets of genes with kinetics and expression patterns unique to circulating or tissue-resident memory CD8^+^ T cells that may contribute to the specification of each distinct subtype. Importantly, these data also revealed transcriptional heterogeneity within the siIEL CD8^+^ T cell population throughout their differentiation. We show that within the established T_RM_ cell population, this molecular heterogeneity reflects functionally distinct subsets that were previously unknown. Moreover, we identify a distinct subset within the CD8^+^ T cell pool within the tissue early in infection that likely represents the precursors of T_RM_ cells. These findings should inform future studies aimed at improving our understanding of T_RM_ cell differentiation and function, which may contribute to our ability to better manipulate CD8^+^ T cell responses to protect against infection and cancer.

## Results

### Single-cell RNA sequencing analyses of circulating and siIEL CD8^+^ T cells responding to viral infection

We employed a single-cell RNA sequencing approach to investigate the transcriptional changes that occur in circulating and siIEL CD8^+^ T cells responding to viral infection. We adoptively transferred P14 CD8^+^CD45.1^+^ T lymphocytes, which have transgenic expression of a T cell receptor (TCR) that recognizes an immunodominant epitope of lymphocytic choriomeningitis virus (LCMV), into congenic CD45.2^+^ wild-type recipients that were infected with the Armstrong strain of LCMV one day later. We FACS-sorted naïve (CD62L^hi^CD44^lo^) P14 T cells from spleens of uninfected mice as well as activated donor P14 T cells (CD44^hi^) from the spleens and siIEL compartments of recipient mice at 11 time points post-infection, and performed scRNA-seq using the 10X Genomics Chromium platform (Fig. 1A and fig. S1).

**Fig. 1.**
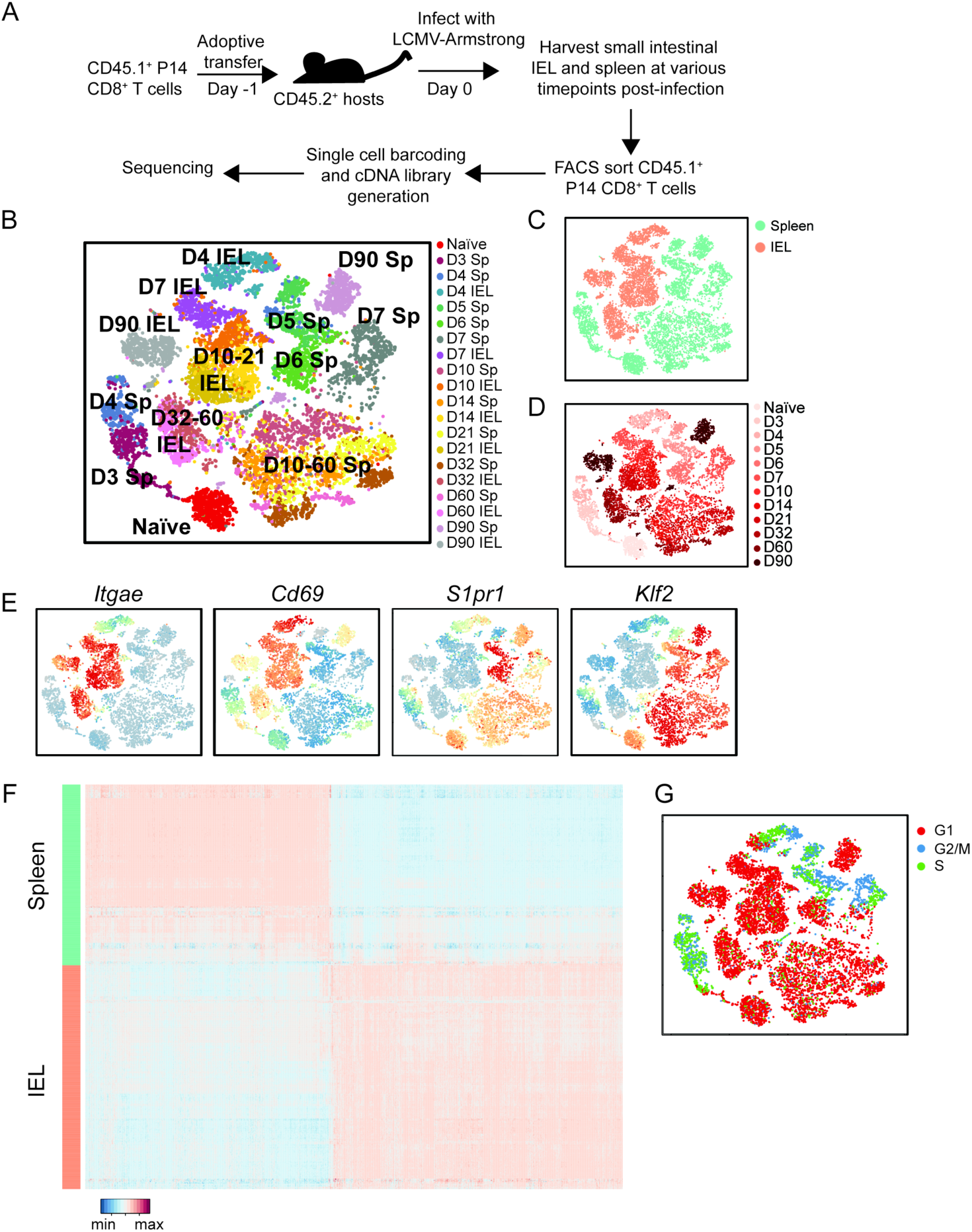
Single-cell RNA sequencing analyses of circulating and siIEL CD8^+^ T cells responding to viral infection. (**A**) Experimental setup. P14 CD45.1^+^CD8^+^ T cells were adoptively transferred into CD45.2^+^ congenic hosts one day prior to infection with LCMV-Armstrong. Splenocytes were harvested at 3, 5, and 6 days; splenocytes and siIEL CD8^+^ T cells were harvested at 4, 7, 10, 14, 21, 32, 60, and 90 days post-infection. Naïve T cells (CD44^lo^CD62L^hi^) were harvested from spleens of uninfected P14 TCR transgenic mice. Antigen-experienced P14 CD8^+^ T cells (CD45.1^+^V*α*2^+^CD44^hi^) were sorted and processed for scRNA-seq with the 10X Genomics platform. (**B** to **D**) tSNE analysis of all scRNA-seq samples, where each individual sample (B), tissue (C), or time point (D) is represented by a unique color. (**E**) Relative expression of known regulators of circulating and tissue-resident memory CD8^+^ T cell differentiation superimposed on individual cells. (**F**) Differential gene expression in Day 4 splenic (teal) and siIEL (coral) CD8^+^ T cells, represented as expression relative to the mean expression among all cells; each row represents an individual cell and each column represents an individual gene. (**G)** Cell cycle status of individual CD8^+^ T cells, inferred from transcriptional profiles.

In order to relate changes in CD8^+^ T cell populations over time and across tissues, we merged the data from all time points and anatomic sites and performed unsupervised t-distributed stochastic neighborhood embedding (tSNE) analysis (Fig. 1, B to D, and fig. S2). CD8^+^ T cells from the siIEL compartment clustered distinctly from splenic CD8^+^ T cells at all time points (Fig. 1 B to D, and fig. S2), consistent with the distinct transcriptional profile of T_RM_ cells that has been previously reported (*14, 15, 18, 20, 21*). For example, expression of genes previously associated with T_RM_ cells, such as *Cd69* and *Itgae,* was strongly enriched among siIEL CD8^+^ T cells compared to splenic CD8^+^ T cells, whereas expression of genes that promote tissue egress and recirculation, such as *Klf2* and *S1pr1*, was significantly lower among siIEL cells compared to splenic cells (Fig. 1E). Strikingly, the divergence in gene-expression profiles between splenic and siIEL CD8^+^ T cells was evident as early as day 4 post-infection (Fig. 1, B to D and fig. S2), the earliest time point at which CD8^+^ T cells can be detected within the intestinal epithelium (*32*), indicating that CD8^+^ T cells begin to change their transcriptional profile rapidly upon entry into tissue. Differential expression analyses revealed that 928 genes were more highly expressed in day 4 splenic CD8^+^ T cells and 1103 genes were more highly expressed in day 4 siIEL CD8^+^ T cells, including *Cd69* and *Itgae* (Fig. 1, E and F and fig. S3, and Table S1). Genes more highly expressed by siIEL CD8^+^ T cells included those associated with processes known to be important for establishment and maintenance of T_RM_ cells, including integrins and cell adhesion molecules (*Itga1*, *Itgb2*, *Itgal*, *Itgb7*, *Itgax*, *Jaml*); regulators of cell trafficking (*Ccr9*, *Cxcr3*) and TGF*β* signaling (*Tgif1*, *Tgfbr2*, *Smad7*, *Skil*, *Smurf2*); the tissue damage receptor *P2xr7*; and fatty acid-binding proteins (*Fabp1*, *Fabp 2*, *Fabp6*) (*33*) (Fig. 1F and fig. S3, and Table S1). Additionally, genes associated with other aspects of T cell function, including cytokines (*Il2*, *Il10*), cytokine receptors (*Il4ra*, *Il2rg*, *Il10rb*, *Il21r*), chemokines (*Ccl3*, *Ccl4*, *Ccl5*, *Cxcl10*, *Cxcl11*), effector molecules (*Gzma*, *Gzmk*, *Fasl*), and both costimulatory (*Icos*, *Cd28*) and inhibitory (*Ctla4*, *Tigit*, *Lag3*) receptors, were more highly expressed by siIEL CD8^+^ T cells. We also observed that genes encoding components and downstream mediators of the TCR signaling pathway (*Zap70*, *Itk*, *Lats2*, *Rasgrp1*, *Fyb*, *Nr4a1*, *Nr4a2*, *Nfatc1*, *Irf4*, *Ikzf2, Cd5*), regulation of intracellular calcium (*Orai1*, *Orai2*, *Sri*, *Rrad*), and regulation of NF-*κ*B signaling (*Nfkbia*, *Nfkbiz*, *Rel*, *Ikbkb*, *Pim1*, *Tnfaip3*) were more highly expressed among siIEL CD8^+^ T cells. Notably, many of these differentially expressed genes were transcription factors with no previously reported role in T_RM_ cells (*Ikzf2*, *Ikzf3*, *Gata3*, *Irf4*, *Id2*). Among the genes more highly expressed by splenic CD8^+^ T cells were *Batf* and *Zeb2,* transcription factors that have been shown to regulate effector CD8^+^ T cell differentiation (*34, 35*); genes encoding for transcription factors, such as *Batf3,* that have not been previously implicated in CD8^+^ T cell differentiation; and genes encoding for the high-affinity IL-2 receptor IL-2R*α* and the costimulatory receptor OX40 (Fig. 1F and fig. S3, and Table S1). Thus, these analyses reveal previously unappreciated signaling pathways and transcription factors that may represent early regulators of circulating versus siIEL CD8^+^ T cell differentiation.

Gene ontology (GO) analyses revealed that genes associated with DNA replication and cell cycle regulation were enriched among splenic CD8^+^ T cells, suggesting that siIEL CD8^+^ T cells may be less proliferative than splenic CD8^+^ T cells (Table S1). Indeed, assessment of cell cycle status inferred from transcriptional analyses suggested that while both splenic and siIEL CD8^+^ T cells were actively dividing at day 4 post-infection, siIEL CD8^+^ T cells had stopped proliferating by day 7 post-infection whereas splenic CD8^+^ T cells continued proliferating until 7-10 days post-infection (Fig. 1G). These findings indicated that siIEL CD8^+^ T cells become quiescent more rapidly following activation compared to splenic CD8^+^ T cells. Taken together, these data reveal that CD8^+^ T cells that enter the siIEL compartment receive signals that alter their transcriptional profile rapidly after arrival, indicating that the T_RM_ cell fate may begin to be specified earlier than previously appreciated, and may serve as a useful resource for identifying early molecular determinants of T_RM_ cell differentiation.

### Shared and tissue-specific components of gene expression programs in circulating and siIEL CD8^+^ T cells

We next sought to elucidate changes in gene-expression programs over time in CD8^+^ T cells responding to viral infection and to understand how gene expression programs in circulating versus siIEL CD8^+^ T cells relate to one another. We analyzed splenic and siIEL CD8^+^ T cells separately and performed weighted gene co-expression network analyses on each set of cells, defining tissue-specific modules of genes that exhibited similar patterns of expression over time in splenic (fig. S4, A and B, and Table S2) or siIEL (fig. S4 C and D, and Table S2) CD8^+^ T cells. We identified 10 and 8 distinct gene-expression modules among splenic and siIEL CD8^+^ T cells, respectively (fig. S4A to D, and Table S2).

Although recent studies have primarily highlighted the differences in gene expression between circulating and tissue-resident CD8^+^ T cells (*14, 15, 18, 20, 21*), we notably found substantial overlap in the gene expression patterns utilized by differentiating splenic and siIEL CD8^+^ T cells. For example, 38% of the genes in Spleen Module 1 were shared with IEL Module 2; genes in these early differentiation modules were characterized by a decrease in expression, relative to that by naïve cells, following T cell activation (fig. S4, B, D, and E). Similarly, 40% of the genes in IEL Module 4 were shared with Spleen Module 4; genes in these intermediate differentiation modules were characterized by high expression during the peak of infection, followed by a subsequent decrease. Lastly, 33% of the genes in IEL Module 7 were shared with Spleen Module 10; genes in these late differentiation modules were characterized by a reduction in expression after T cell activation, followed by increasing expression that was sustained at late time points (fig. S4, B, D, and E, and Table S2). These observations suggested that splenic and siIEL CD8^+^ T cells share a core transcriptional program throughout their differentiation, consistent with their common cytolytic lymphocyte functions (fig. S4E and Table S2), and highlight the potential value of this resource dataset to reveal how specific genes or pathways, including those known to play key roles in regulating circulating CD8^+^ T cell memory, may act similarly or disparately in regulating tissue-resident memory differentiation.

To elucidate distinct regulators of the circulating and tissue-resident memory differentiation programs, we again performed weighted gene co-expression network analyses, but this time analyzed splenic and siIEL CD8^+^ T cells together. This approach enabled us to directly compare the level of representation of each module in cells from each anatomic site at each time point. Fifteen modules representing distinct temporal patterns of gene expression were defined and annotated as Combined Modules 1 – 15 (Fig. 2, A to C, and Table S3). This analysis confirmed our observation that splenic and siIEL CD8^+^ T cells indeed share a core transcriptional program, as many Combined Modules were similarly represented in splenic and siIEL CD8^+^ T cells throughout most of their differentiation (Fig. 2, B and C). Moreover, this analysis confirmed our finding that siIEL CD8^+^ T cells were transcriptionally distinct from splenic CD8^+^ T cells as early as day 4 post-infection (Fig. 1, B to F), since we noted that many modules (Combined Modules 2, 3, 5, 7, 11, and 12) were differentially represented within splenic and siIEL CD8^+^ T cells at day 4 post-infection, but not at later time points (Fig. 2, B and C). For example, genes in one such module, Combined Module 12, were enriched in several pathways known to play roles in circulating CD8^+^ T cell memory, including autophagy, protein ubiquitination, and proteasome protein catabolism (*36, 37*) (Table S3). These findings suggested that differentiating siIEL CD8^+^ T cells may begin to acquire certain aspects of a memory-like transcriptional profile more rapidly than splenic CD8^+^ T cells.

**Fig. 2.**
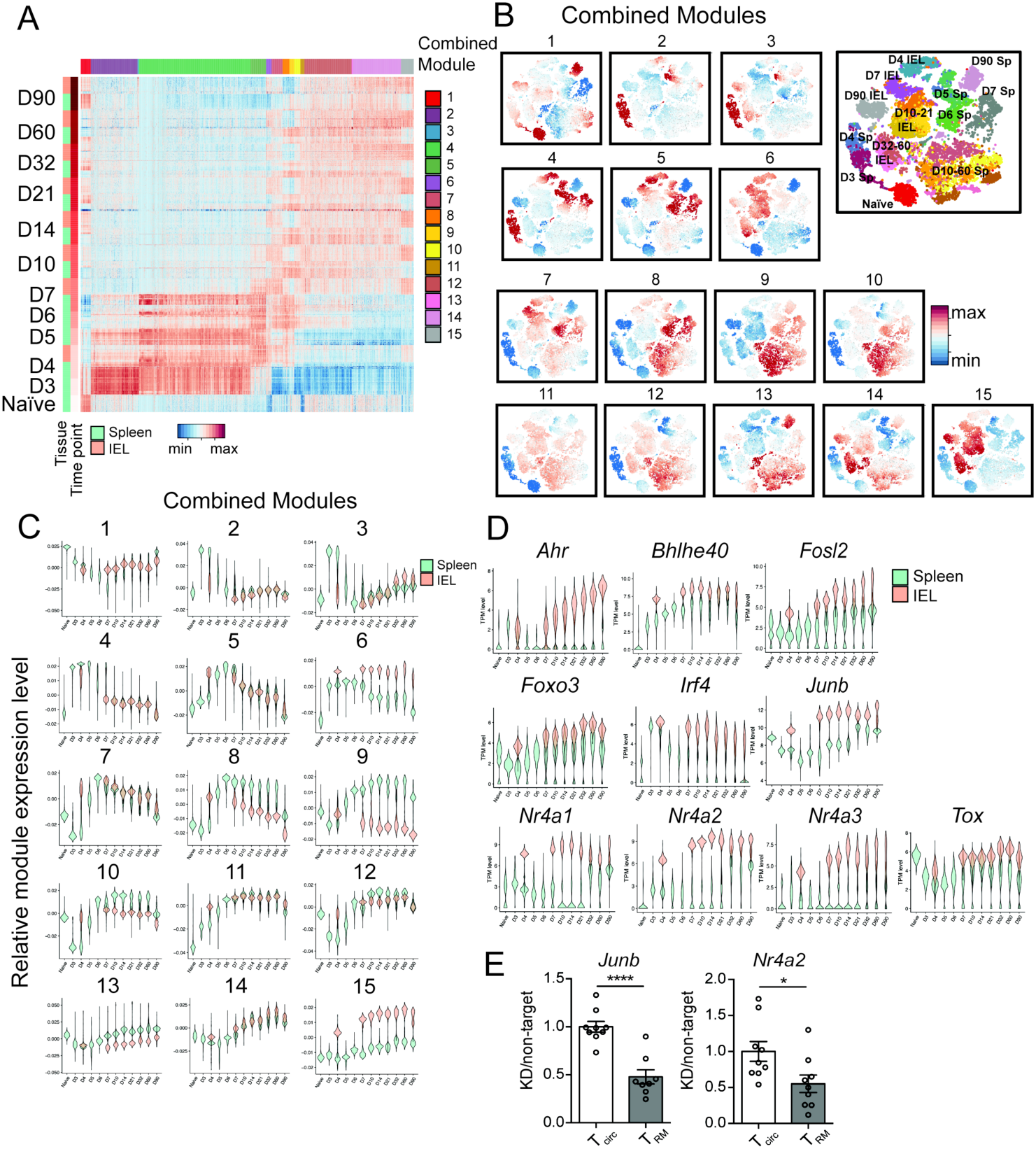
Identification of a siIEL CD8^+^ T cell-enriched gene-expression profile. (**A**) Gene-expression patterns of single splenic and siIEL CD8^+^ T cells responding to infection, where each row represents an individual cell, grouped by tissue within each time point, and each column represents an individual gene, grouped by module, represented as expression relative to the mean expression among all cells. Weighted gene co-expression network analyses of splenic and siIEL CD8^+^ T cells considered together were performed to derive gene modules. (**B** and **C**) Representation of each module among single CD8^+^ T cells in both the spleen and siIEL compartment over time, relative to the mean representation among all cells, depicted as a representation score superimposed on individual cells (B) or as violin plots showing the relative representation of each module in individual splenic (teal) versus siIEL CD8^+^ T cells (coral) (C). (**D**) Violin plots depicting gene-expression patterns of known or putative regulators of T_RM_ cell differentiation (represented as transcripts per million, TPM), selected from siIEL CD8^+^ T cell-enriched modules, among single splenic (teal) or siIEL (coral) CD8^+^ T cells over time. (**E**) CD45.1^+^ P14 T cells were transduced with retrovirus encoding shRNA targeting *Nr4a2* or *Junb* (knockdown, KD), and mixed with CD45.1.2^+^ P14 T cells transduced with shRNA encoding control (non-target) shRNA at a 1:1 ratio of KD: non-target cells prior to adoptive transfer into CD45.2^+^ hosts that were subsequently infected with LCMV. 22-26 days later, splenic and siIEL CD8^+^ T cells were analyzed by flow cytometry. Ratio of KD: non-target P14 T cells within transduced P14 CD8^+^ T cells in the spleen (T_circ_) or within transduced CD69^+^CD103^+^ siIEL P14 CD8^+^ T cells (T_RM_), normalized to the ratio of KD: non-target cells at the time of transfer. Values are normalized to T_circ_. *P<0.05, ****P<0.0001 (Student’s two-tailed *t*-test). Data were pooled from two independent experiments, with mean and SEM of n=8 mice per gene, where each dot represents an individual mouse.

These analyses also defined several modules that were significantly and consistently differentially enriched in splenic vs. siIEL CD8^+^ T cells over time. For example, Combined Modules 6 and 15 were enriched in siIEL compared to splenic CD8^+^ T cells; genes in these modules were highly upregulated in siIEL CD8^+^ T cells throughout their differentiation (Figures 2B and 2C). Combined Modules 6 and 15 contained 157 and 371 genes, respectively, some of which have been previously implicated in CD8^+^ T_RM_ cell differentiation and function, such as *Cd69*, *Itgae*, *Itga1*, and *Ccr9*, which regulate homing and retention within the intestinal environment; transcription factors *Nr4a1* and *Ahr*, which regulate the long-term maintenance of T_RM_ (*38, 39*), and *Runx3*, a transcriptional regulator of key genes that establish tissue-residency that is critical for T_RM_ cell differentiation (*18*); and *Ccl3* and *Ccl4*, which are involved in the rapid recruitment of innate cells by T_RM_ cells (*11, 12*) (Fig. 2D and fig. S5, Table S3). Additionally, GO analyses revealed enrichment of pathways known to play roles in T_RM_ cell differentiation and maintenance, such as TGF*β* signaling (*17*) (*Skil*, *Smurf, Tgif1*) and fatty acid homeostasis (*Dgat1*, *Got1*), consistent with the dependence of T_RM_ cells on fatty acid metabolism for their maintenance and function (*33*) (Fig. 2D and fig. S5, and Table S3). These T_RM_ cell-enriched modules also included genes encoding effector molecules and cytokines, such as *Il2*, *Ifng, Tnf, Fasl,* and *Gzmb*, sustained high expression of which may contribute to the rapid and potent recall responses of T_RM_ cells. Moreover, other groups of genes represented in these modules implicated pathways with previously unexplored roles in T_RM_ cell differentiation, some of which were already upregulated in siIEL CD8^+^ T cells as early as day 4 post-infection (Fig. 1F and fig. S3). These included inhibitory receptors (*Ctla4*, *Lag3*, *Cd101*, *Tigit*); factors involved in TCR signaling and costimulation (*Zap70*, *Lat*, *Cd8a*, *Cd3e, Cd40lg, Nfatc1*); regulators of cell survival (*Bcl2a1b*, *Bcl2l11*, *Bcl2a1d*) and NF-*κ*B signaling (*Nfkbia*, *Nfkbiz*, *Nfkbid*, *Rel*); genes involved in cytokine responses (*Stat3*, *Stat1*, *Irak2*, *Socs1*, *Jak2*); and transcription factors with previously uncharacterized roles in T_RM_ cell differentiation (*Nr4a2*, *Nr4a3*, *Junb*, *Fosl2*, *Tox*, *Bhlhe40*, *Foxo3, Irf4*) (Fig. 2D and fig. S5, and Table S3). In addition, genes involved in cholesterol biosynthesis (*Fdps*, *Cyp51*, *Mvk*, *Fdft1*, *Hmgcr*, *Hmgcs*) and steroid hormone mediated signaling pathways were enriched in Combined Modules 6 and 15, respectively, raising the possibility of a role for these processes in T_RM_ cell differentiation.

By contrast, Combined Modules 8 and 9 were enriched in splenic relative to siIEL CD8^+^ T cells beginning at day 7 post-infection, and included 196 and 138 genes, respectively, some of which have been previously implicated in promoting differentiation of circulating memory CD8^+^ T cells at the expense of T_RM_ cells (Fig. 2, B and C, and Table S3). For example, these spleen-enriched modules included *Klf2*, *S1pr1*, *Ly6c1,* and *Ly6c2*, which regulate trafficking and egress, and *Eomes*, a transcription factor that negatively regulates T_RM_ cell differentiation (*14, 16, 17, 32*) (fig. S5 and Table S3). These spleen-enriched modules also included several cell surface markers, including *Cx3cr1* and *Klrg1,* that have been associated with subsets of circulating effector CD8^+^ T cells. Additionally, the presence of genes encoding specific cytokine receptors (*Il18r1*, *Il18rap*, *Il17ra*) in these spleen-enriched modules suggests that signaling through these receptors might negatively regulate T_RM_ cell differentiation and maintenance.

In order to demonstrate that genes identified by these analyses indeed represent functionally important regulators of T_RM_ cell differentiation, we selected for further studies two genes found in Combined Module 15, Nuclear Receptor Subfamily 4 Group A Member 2 (*Nr4a2*) and Junb proto-oncogene, AP-1 transcription factor subunit (*Junb*), both of which exhibited increased expression in siIEL relative to splenic CD8^+^ T cells at all time points (Fig. 2D). *Nr4a2* is an orphan nuclear receptor that belongs to the Nr4a family of transcription factors, plays a central role in the development of regulatory T cells as well as pathogenic Th17 cells (*40–43*), and has been implicated as an important driver of exhaustion in CD8^+^ T cells (*44*). *Junb* plays important roles in regulating Th17 cell identity and pathogenicity (*45, 46*), and has also been implicated as part of the effector CD8^+^ T cell transcriptional program (*35*). Neither *Nr4a2* nor *Junb* have previously reported roles in the differentiation of T_RM_ cells. To investigate whether these genes play a role in T_RM_ cell differentiation, we transduced P14 CD8^+^CD45.1^+^ T cells with retroviruses encoding shRNA targeting *Nr4a2* or *Junb*, and transduced P14 CD8^+^CD45.1.2^+^ T cells with retroviruses encoding non-targeting shRNA. Cells were mixed at a 1:1 ratio and adoptively transferred into CD45.2 recipient mice that were subsequently infected with LCMV (fig. S6A). Relative to non-targeting controls, knockdown of *Nr4a2* or *Junb* resulted in a greater reduction in siIEL T_RM_ cells than splenic memory CD8^+^ T cells (Fig. 2E and fig. S6B). Taken together, these results highlight key transcriptional differences between circulating and siIEL CD8^+^ T cells undergoing differentiation and demonstrate the potential value of this resource dataset in identifying previously unknown genes and pathways involved in specifying T_RM_ versus circulating memory CD8^+^ T cell fates.

### Single-cell RNA-sequencing analyses highlight circulating CD8^+^ T cell heterogeneity and reveal previously unappreciated heterogeneity within the siIEL CD8^+^ T cell pool

We next investigated potential heterogeneity of differentiating splenic and siIEL CD8^+^ T cells. We observed that individual splenic CD8^+^ T cells at certain time points expressed genes previously associated with CD8^+^ T cell subsets, such as terminal effector (TE)-and memory precursor (MP)-phenotype cells at the peak of infection, and T_EM_, T_CM_, and LLE cell subsets at later time points following infection (Fig. 3, A and B). For example, the majority of cells at days 6 and 7 post-infection expressed high levels of *Klrg1*, likely representing TE cells, whereas a smaller number of cells exhibited lower expression of *Klrg1* and higher expression of *Tcf7* and *Bcl2*, likely representing MP cells (*47–49*). We also observed that certain cells at later time points expressed genes previously associated with T_CM_ (*Il7r*, *Tcf7*, *Sell, Bcl2, Cxcr3*), LLE (*Klrg1*, *Cx3cr1*, *Zeb2*), and T_EM_ (intermediate levels of *Cx3cr1*, *Zeb2*, *Il7r*, *Tcf7,* and *Bcl2*) cells (*1, 2*) (Fig. 3, A and B). Indeed, mapping individual splenic CD8^+^ T cells to TE, MP, T_CM_, T_EM_, and LLE transcriptional signatures defined by bulk RNA-seq signatures of sorted cells from each subset, revealed that heterogeneity observed among splenic CD8^+^ T cells might indeed represent cells differentiating into each of these memory cell subsets (Fig. 3, C and D). Notably, low, intermediate, or high expression of *Cx3cr1* at day 6 post-infection generally corresponded to cells with T_CM_, T_EM_, or LLE profiles, respectively, consistent with previously reported CX3CR1^hi^, CX3CR1-intermediate, and CX3CR1^lo^ subsets with distinct functional capacities and terminal differentiation potentials that can be found at the peak of infection(*7*) (Fig. 3, B and D). These results demonstrate the concurrent presence of these memory subsets within the circulating CD8^+^ T cell memory pool, consistent with previous studies (*4–6, 8*), and support findings suggesting that these memory CD8^+^ T cell subsets may begin to diverge in their differentiation pathways earlier than is widely appreciated (*7*). Moreover, our dataset might provide a useful resource for identifying previously undescribed regulators of circulating CD8^+^ T cell heterogeneity and function.

**Fig. 3.**
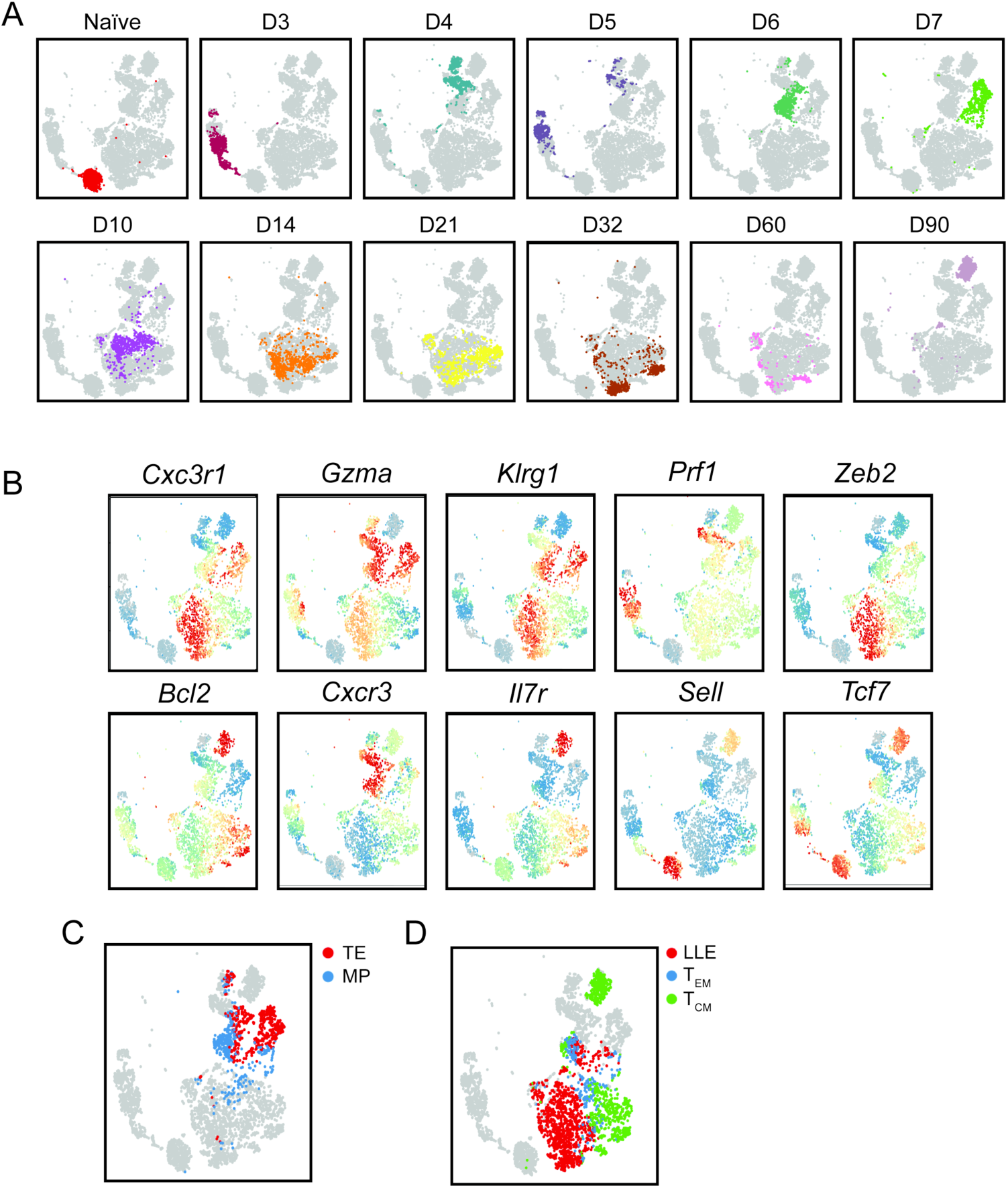
Single-cell RNA sequencing analyses highlight heterogeneity among circulating CD8^+^ T cells. (**A**) tSNE analysis of splenic CD8^+^ T cells; each plot represents an individual time point with each color representing a specific time point, as in Fig. 1B. (**B**) Relative expression (compared to mean expression among all spleen cells) of known regulators of circulating CD8^+^ T cell differentiation superimposed on individual spleen cells. (**C** and **D**) Similarity of gene-expression programs among single spleen cells to transcriptional signatures (derived from bulk RNA-seq profiles from FACS-sorted cells) of previously defined circulating CD8^+^ T cell subsets. (C) Similarity of gene-expression programs among single spleen cells to terminal effector (TE) (D7 KLRG1^hi^IL-7R*α*^lo^, red) or memory-precursor (MP) (D7 KLRG1^lo^IL-7R*α*^hi^, blue) transcriptional signatures. (D) Similarity of gene-expression programs among single spleen cells to LLE (D35 CD62L^lo^IL-7R*α*^lo^, red), T_EM_ (D35 CD62L^lo^IL-7R*α*^hi^, blue), and T_CM_ (D35 CD62L^hi^IL-7R*α*^hi^, green) cell transcriptional signatures.

Although heterogeneity within circulating CD8^+^ T cells has been relatively well described, whether heterogeneity exists within tissue-resident CD8^+^ T cell populations remains unclear. To investigate potential heterogeneity within the siIEL CD8^+^ T cell compartment, we defined cell clusters across the entire dataset, revealing 40 distinct clusters, 17 of which were found in the siIEL compartment (Fig. 4A). We observed that multiple clusters of siIEL CD8^+^ T cells were present within several individual time points, suggesting that several distinct subsets might exist within the siIEL CD8^+^ T cell population (Fig. 4B). For example, at day 60 post-infection, two major cell clusters, Clusters 3 and 29, were observed. These clusters exhibited differential expression of 236 genes, including several transcription factors, suggesting that these clusters were regulated by distinct gene-expression programs (Table S4). For example, Cluster 3 cells had higher expression of *Id3, Jun, Fos, Klf2,* and *Myc*, while Cluster 29 cells had higher expression of transcription factors including *Bcl6, Zeb2*, and *Klf3*. Cluster 3 cells also exhibited higher expression of several genes associated with T_RM_ cell function, including cytokines and chemokines (*Ifng*, *Tnf*, *Ccl3*, *Ccl4*), suggesting that Cluster 3 cells might be better poised to produce cytokines rapidly upon re-infection (Fig. 4C, Table S4). These analyses suggest that distinct costimulatory and other signaling pathways might be responsible for regulating each of these cell clusters and raised the possibility that these cell clusters might have distinct functional capacities.

**Fig. 4.**
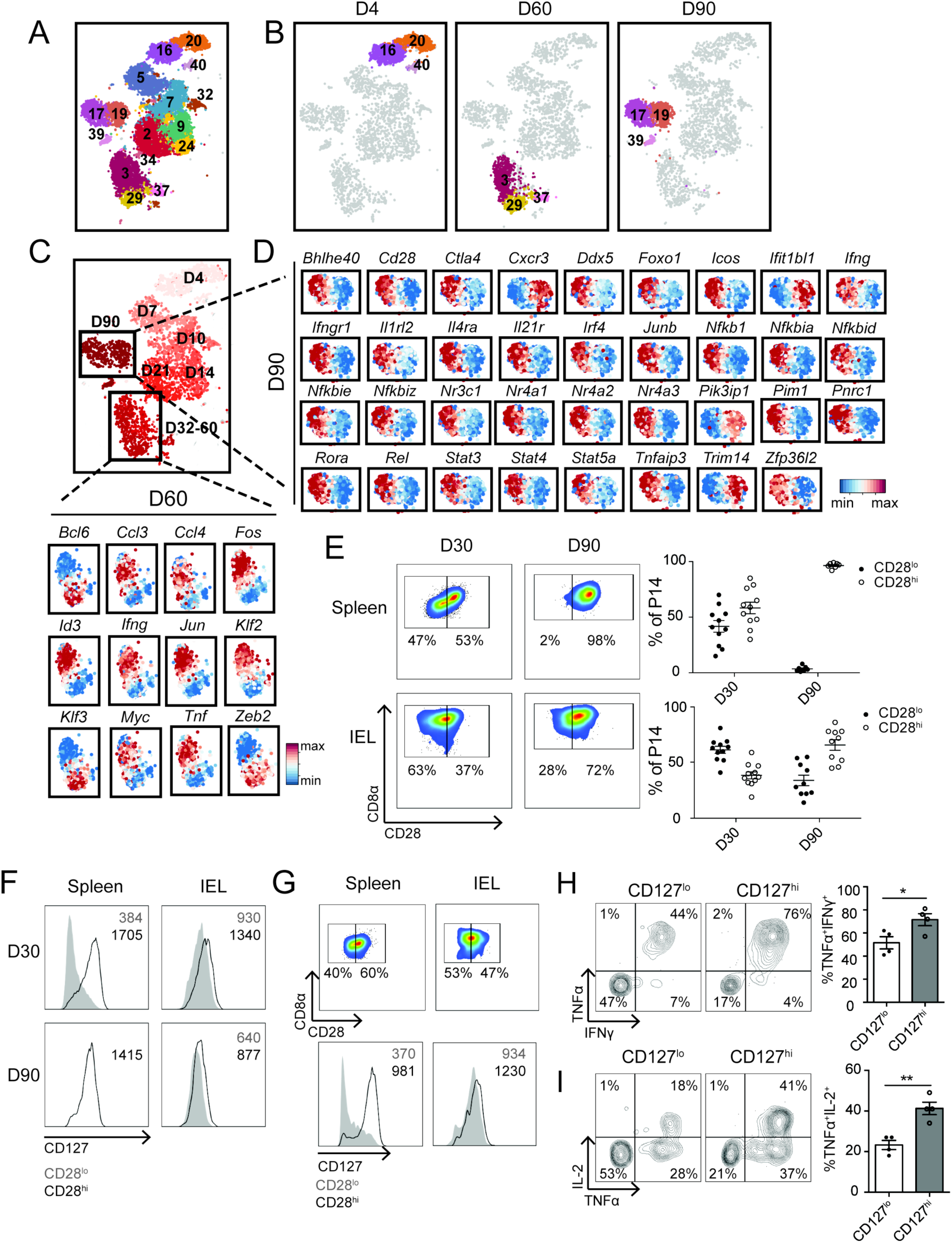
Single-cell RNA sequencing analyses reveal functionally distinct subsets within the siIEL CD8^+^ T cell pool. (**A** and **B**) Clustering analyses of siIEL CD8^+^ T cells from all time points (A) or cells at days 4, 60, and 90 post-infection (B); numbers represent different cluster annotations. (**C** and **D**) Expression of selected genes within single siIEL CD8^+^ T cells at day 60 (C) or 90 (D) post-infection, relative to the mean expression among siIEL CD8^+^ T cells at the indicated time point, demonstrating differential gene expression between Clusters 3 and 29 (C) or between Clusters 17 and 19 (D). (**E** to **I**) P14 CD8^+^ T cells were transferred into congenically distinct recipients 1 day prior to infection with LCMV. Splenic and siIEL CD8^+^ T cells were harvested at day 30 or 90 post-infection. (E) Representative flow cytometry plots displaying distribution of CD28 expression among total P14 T cells in the spleen (top) or CD69^+^CD103^+^ P14 T cells in the siIEL compartment (bottom). Numbers represent the percentage of total P14 T cells in each gate, and graphs to the right demonstrate the quantification of CD28^lo^ (filled dots) and CD28^hi^ (open circles) cells within the indicated population at each time point. Data are representative of (flow cytometry plots, left), or compiled from (graphs, right) 3 independent experiments where n=2-5 mice per experiment (10-11 total mice). (F) Expression of CD127 (IL-7R*α*) by CD28^lo^ (gray) or CD28^hi^ (black) subsets within the spleen (left) or siIEL (right) at day 30 (top) and 90 (bottom) post-infection. Numbers represent Mean Fluoresence Intensity (MFI) of CD127 within the indicated population. (G) Representative flow cytometry plots displaying (top) distribution of CD28 expression among total H-2D^b^ GP_33-41_ tetramer^+^ endogenous siIEL CD8^+^ T cells in the spleen (left) or H-2D^b^ GP_33-41_ tetramer^+^ endogenous CD69^+^CD103^+^ CD8^+^ T cells in the siIEL compartment (right), or displaying (bottom) expression of CD127 by CD28^lo^ (gray) or CD28^hi^ (black) subsets within the spleen (left) or siIEL (right). Numbers represent the percentage of H-2D^b^ GP_33-41_ tetramer^+^ cells in each gate (top), or MFI of CD127 expression in the indicated population (bottom) (H-I) At day 30 post-infection, siIEL P14 T cells were sorted into CD127^lo^ and CD127^hi^ subsets and cultured in the presence of GP_33-41_ peptide *in vitro*. Quantification (right) of the proportion of CD127^hi^ or CD127^lo^ P14 siIEL CD8^+^ T cells producing IFN*γ* and TNF*α* (H), or IL-2 and TNF*α* (I) as shown in representative flow cytometry plots (left). Data are representative of 2 independent experiments, with mean and SEM of n=4 wells of cultured cells per phenotype, derived from two separate pools of sorted cells plated in duplicate.

Additionally, at day 90 post-infection, two major cell clusters, Clusters 17 and 19, were observed; these clusters exhibited differential expression of 361 genes (Table S5). Cluster 17 cells exhibited higher expression of 243 genes, including transcription factors (*Bhlhe40, Ddx5, Foxo1, Irf4, Junb, Nr3c1, Nr4a1*, *Nr4a2*, *Nr4a3, Pnrc1,* and *Rora*) and mediators of cytokine signaling (*Ifngr1, Il1rl2 Il21r, Il4ra,* and *Stat3, Stat5a,* and *Stat4*). Cluster 17 cells also exhibited higher expression of *Zfp36l2*, which encodes for a zinc finger protein involved in the regulation of mRNA decay, and the transcript for IFN*γ*, as well as genes associated with pathways that were highly represented within T_RM_ cell-enriched modules, such as the regulation of NF-*κ*B signaling (*Nfkbia, Nfkbid, Tnfaip3, Pim1, Rel, Nfkb1, Nfkbie, Nfkbiz*) and costimulatory and inhibitory molecules *Icos* and *Ctla4* (Fig. 4D and Table S5). Cluster 17 cells also exhibited higher expression of the costimulatory receptor *Cd28*, suggesting that costimulation through this receptor could specifically influence the differentiation or responsiveness of a specific subset of T_RM_ cells. By contrast, Cluster 19 exhibited higher expression of 117 genes, including the tissue-homing molecule *Cxcr3*, interferon response genes (*Trim14, Ifit1bl1*), and *Pik3ip1*, a negative regulator of T cell activation (Fig. 4D and Table S5). Taken together, these findings suggested that specific cytokine and/or costimulatory signals might be responsible for maintaining or directing the differentiation of Cluster 17 cells. Moreover, while Cluster 17 cells might be better poised to produce cytokines rapidly upon re-infection, Cluster 19 cells might be more likely to respond immediately to type I interferon signals within the tissue. These data suggest that multiple subsets with distinct transcriptional programs and functional capacities may exist within the T_RM_ cell pool.

We next performed flow cytometry to determine whether the heterogeneity revealed by scRNA-seq analyses could be discerned at the protein level. P14 CD8^+^ T cells were adoptively transferred into congenically distinct hosts that were infected with LCMV one day later. At days 30 and 90 post-infection, the spleens and siIEL compartments of recipient mice were harvested for analysis. At day 30 post-infection, both splenic and siIEL CD8^+^ T cells could be divided into CD28^hi^ and CD28^lo^ populations (Fig. 4E). In contrast, at day 90 post-infection, splenic P14 CD8^+^ T cells uniformly expressed high levels of CD28, but heterogeneity was retained within the siEL P14 CD8^+^ T cell population (Fig. 4E), in line with data revealing multiple clusters within the T_RM_ cell population with differential expression of *Cd28* at the mRNA level at day 90 post-infection (Fig. 4D). We also noted that expression of IL-7R*α* (CD127) also correlated with expression of CD28 (Fig. 4F). Notably, we also observed this heterogeneity among endogenous, siIEL CD8^+^ H-2D^b^ GP_33-41_ tetramer^+^ cells, with CD28^hi^ cells expressing higher levels of CD127 (Fig. 4G). To investigate whether heterogeneity in expression of these markers might indeed reflect functional heterogeneity, we sorted CD127^hi^ and CD127^lo^ populations from the siIEL CD8^+^ T cell pool at day 30 post-infection, and measured their capacities to produce cytokines when rechallenged with cognate antigen *in vitro*. We found that CD127^hi^ cells produced higher levels of the cytokines IFN*γ*, TNF*α*, and IL-2 in response to restimulation than did CD127^lo^ cells (Fig. 4, H and I), consistent with the higher level of *Ifng* transcript detected within CD28^hi^ cells at day 90 post-infection (Fig. 4, C and D). Taken together, these data demonstrate that the heterogeneity revealed by scRNA-seq analyses within the T_RM_ cell pool at late time points after infection is functionally important.

The observation that a number of transcriptional regulators, including *Ddx5, Junb*, *Nr4a2*, and *Pnrc1*, as well as the RNA-binding protein *Zfp36l2*, correlated with *Cd28* expression (Fig. 4D), raised the possibility that these genes might represent regulators of T_RM_ cell heterogeneity. To investigate this possibility, we transduced P14 CD8^+^CD45.1^+^ T cells with retroviruses encoding shRNA targeting *Junb*, *Nr4a2*, *Pnrc1*, or *Zfp36l2* prior to adoptive transfer into congenically distinct hosts that were subsequently infected with LCMV. Knockdown of these genes resulted in a reduction in the proportion of siIEL T_RM_ cells expressing high levels of CD28 compared to congenically distinct, co-transferred control P14 CD8^+^ T cells transduced with retroviruses encoding non-targeting shRNA (Fig. 5A and fig. S6A). Additionally, we co-transferred P14 CD8^+^ T cells from mice with a T cell-specific deletion of *Ddx5* (*Ddx5^fl/fl^*CD4cre^+^: ‘*Ddx5*^-/-^’) and congenically distinct control P14 CD8^+^ T cells (*Ddx5^fl/fl^*CD4cre^-^: ‘wild-type, WT’) into recipients infected with LCMV one day later, and found that the proportion of CD28^hi^ cells in the siIEL compartment was reduced among *Ddx5*^-/-^ cells (Fig. 5B). Moreover, consistent with this decrease in the proportion of CD28^hi^ cells, *Ddx5*^-/-^ siIEL, but not splenic, CD8^+^ T cells, exhibited a reduced ability to produce IFN*γ* and TNF*α* upon restimulation with cognate antigen *in vitro* (Fig. 5C). Taken together, these findings suggest that *Ddx5*, *Junb*, *Nr4a2*, *Pnrc1*, and *Zfp36l2* regulate the differentiation of transcriptionally and functionally distinct T_RM_ cell subsets, and demonstrate the potential value of this resource dataset to reveal regulators of functional heterogeneity within the T_RM_ cell population.

**Fig. 5.**
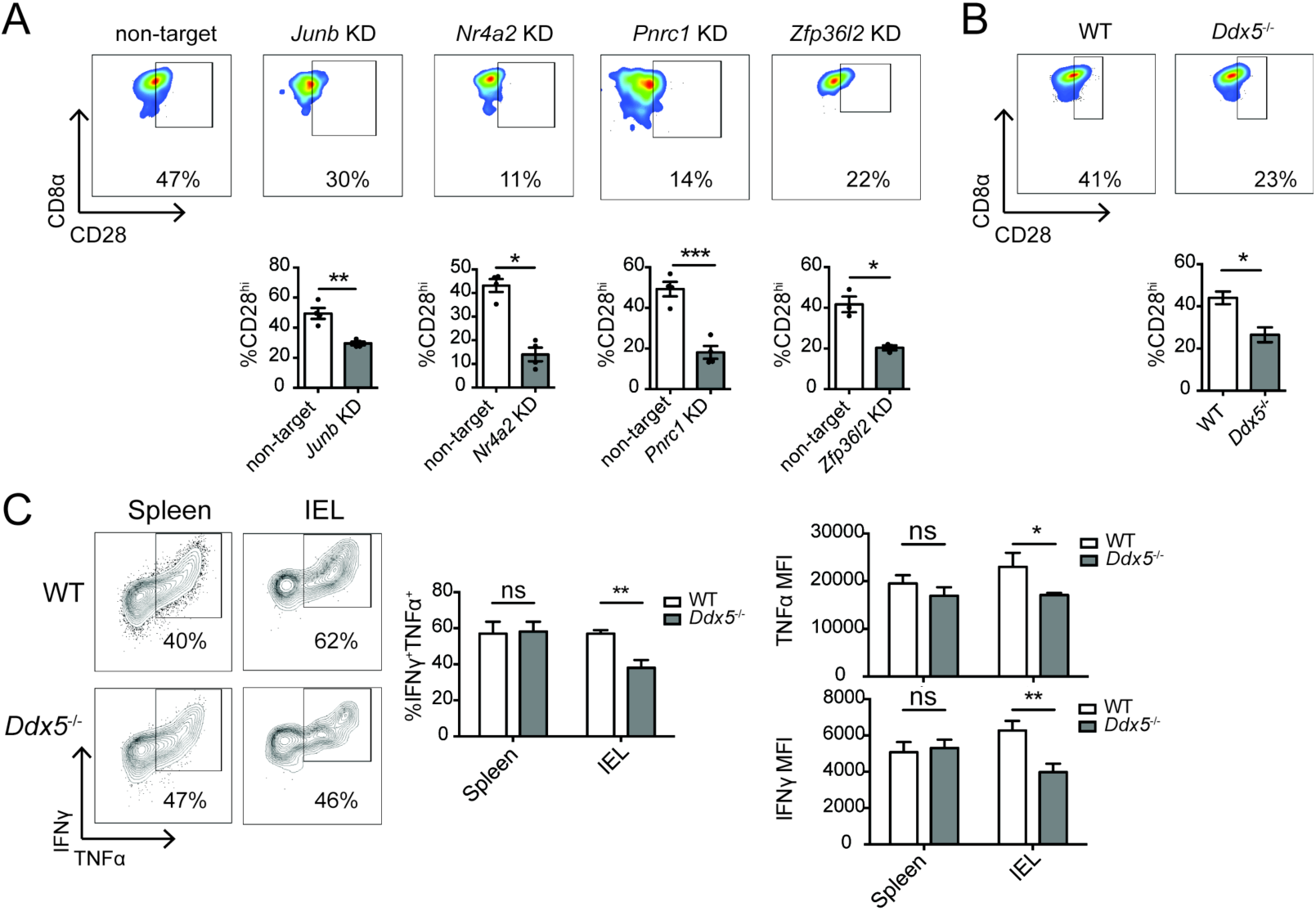
Single-cell RNA sequencing analyses identify putative regulators of CD8^+^ T_RM_ cell heterogeneity. (**A**) CD45.1^+^ P14 T cells were transduced with retrovirus encoding shRNA targeting the indicated genes (knockdown, KD), and mixed with CD45.1.2^+^ P14 T cells transduced with shRNA encoding control (non-target) shRNA at a 1:1 ratio of KD: non-target cells prior to adoptive transfer into CD45.2^+^ hosts that were subsequently infected with LCMV, as in Figure S6. 22-23 days later, siIEL CD8^+^ T cells were analyzed by flow cytometry. Quantification of the percent of non-target or KD P14 siIEL CD8^+^ T cells expressing high levels of CD28 (bottom) as shown in representative flow cytometry plots (top). Data are representative of 1-2 experiments, with mean and SEM of n=4 mice per gene, where each dot represents an individual mouse. (**B** and **C**) P14 CD8^+^ T cells from mice with T cell-specific deletion of *Ddx5* (*Ddx5^fl/fl^* CD4-Cre^+^: ‘*Ddx5*^-/-^’) were co-transferred at a 1:1 ratio with congenically distinct control P14 CD8^+^ T cells (*Ddx5^fl/fl^* CD4-Cre^-^: ‘WT’) into congenically distinct hosts that were infected with LCMV one day later. At day 30 post-infection, spleens and siIEL were harvested for flow cytometric analysis. (B) Quantification of the percent of *Ddx5*^-/-^ or WT siIEL P14 CD8^+^ T cells expressing high levels of CD28 (bottom) as shown in representative flow cytometry plots (top). (C) Lymphocytes harvested from the spleen and small intestinal epithelium were cultured in the presence of GP_33-41_ peptide *in vitro* prior to staining for flow cytometric analysis. Cytokine production in *Ddx5*^-/-^ or WT siIEL P14 CD8^+^ T cells, shown as quantification of the level of TNF*α* or IFN*γ* produced, displayed as MFI of TNF*α* within TNF*α*^+^ cells, or MFI of IFN*γ* within IFN*γ*^n^ cells, or the proportion of cells producing TNF*α* and IFN*γ*, as shown in representative flow cytometry plots (left). Data are quantified from 1 experiment with mean and SEM of n=3 mice per genotype. *P<0.05, **P<0.01, ***P<0.001 (Student’s two-tailed *t*-test).

Although it is known that two distinct populations with differing memory potentials, distinguished by high or low expression of the high-affinity IL-2 receptor, IL-2R*α* (CD25) {Arsenio:2014jx, Kalia:2010hn}, exist within the circulating CD8^+^ T cell pool early in infection, whether the CD8^+^ T cell pool that seeds the tissue early in infection is heterogeneous remained unknown. To determine whether heterogeneity of siIEL CD8^+^ T cells can be discerned early after infection, we next examined the earliest time point, day 4 post-infection, at which CD8^+^ T cells could be detected in the siIEL compartment. In addition to the heterogeneity observed at late time points following infection (Fig. 4B), two major clusters, Clusters 16 and 20, were evident at day 4 post-infection and exhibited differential expression of 332 genes (Fig. 4B, 5B and 6A, and Table S6). Only 17 of these genes were more highly expressed by Cluster 16 cells, such as interferon response genes *Ifit1*, *Ifit1bl1,* and *Ifit3*; *Pik3ip1*, a negative regulator of TCR signaling; *Mxd4*, a Myc-antagonist that promotes survival in T cells(*50*); and *Kdm5b*, a histone demethylase (Fig. 6A and Table S6). Gene Ontology analyses revealed that the genes expressed more highly by Cluster 20 cells were overrepresented in biologic processes such as DNA replication, mitotic spindle and nucleosome assembly, cytokinesis, and cell division; specific genes included *Cdc7*, *Cdc25b*, *Cenpk*, *Cenpm*, *Cdc20*, and *Cdk1* (Fig. 6A and Table S6). Moreover, additional analyses confirmed that a greater proportion of Cluster 20 cells were in the G2/M phases of the cell cycle compared to Cluster 16 cells, indicating that cells in Cluster 20 were more actively proliferating (Fig. 6B). Cluster 20 cells also expressed higher levels of *Dnmt1*, a DNA methyltransferase that is critical for the expansion of CD8^+^ T cells during the effector phase of the response (*51*), along with several components of the Polycomb Repressive Complex 2, including *Suz12* and *Ezh2* (Fig. 6A), which negatively regulates gene expression via histone methylation and has been previously shown to promote effector CD8^+^ T cell differentiation by mediating the repression of memory-associated genes (*22, 52*). These results suggested that two distinct populations of CD8^+^ T cells are present within the siIEL compartment at 4 days post-infection.

**Fig. 6.**
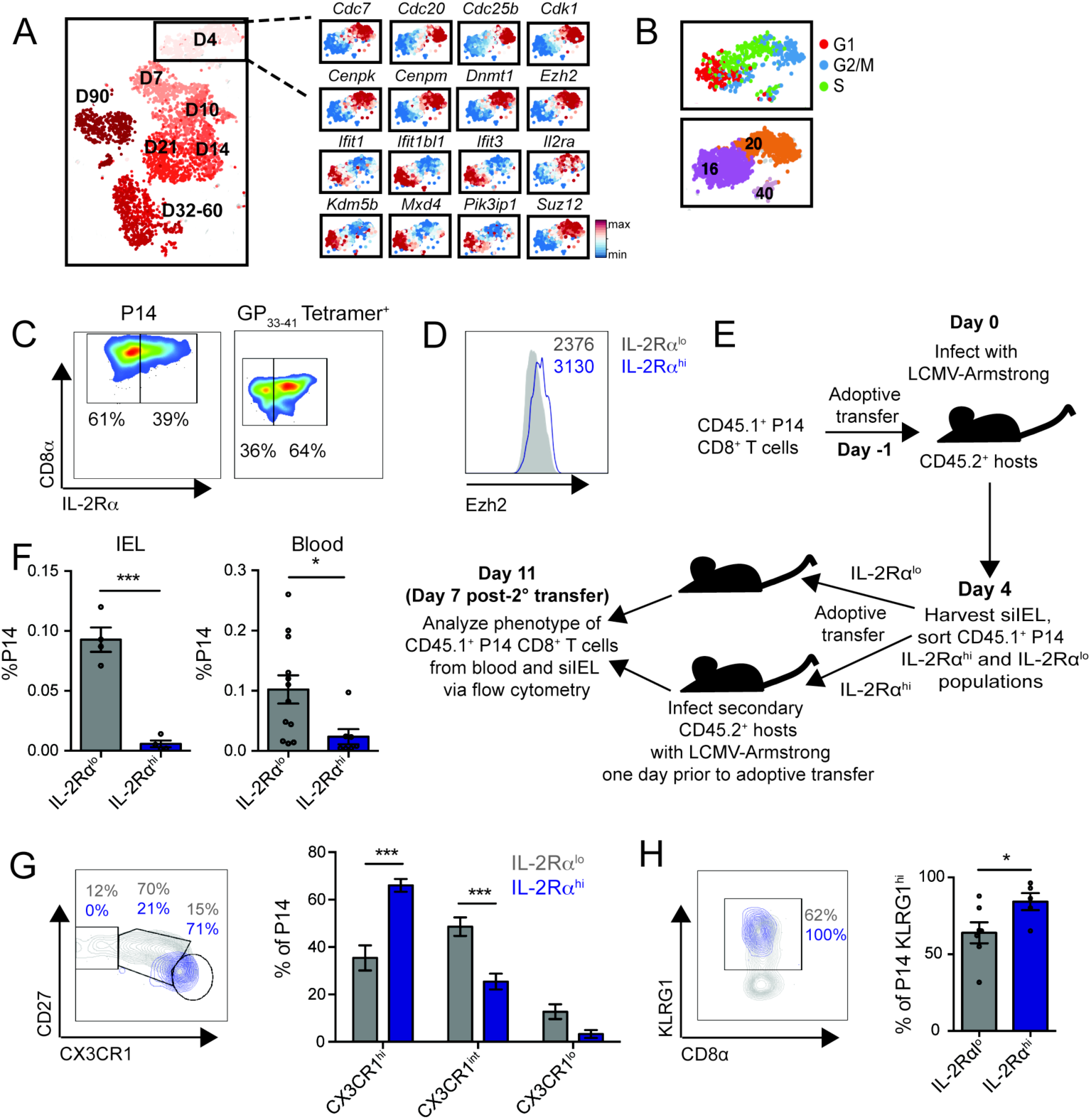
T_RM_ precursors identified within the siIEL CD8^+^ T cell pool early in infection. (**A**) Expression of selected genes within single siIEL CD8^+^ T cells at day 4 post-infection, relative to the mean expression among all cells, demonstrating differential gene expression between cells in Clusters 16 and 20. (**B**) Cell cycle status of siIEL CD8^+^ T cells at day 4 post-infection, inferred from transcriptional profiles. (**C** and **D**) P14 CD8^+^ T cells were adoptively transferred into congenic hosts one day prior to infection with LCMV. Splenic and siIEL CD8^+^ T cells were analyzed by flow cytometry at day 4 post-infection. Data are representative of two independent experiments, with n=2-3 mice per experiment. (C) Representative flow cytometry plots displaying distribution of IL-2R*α* expression among total siIEL P14 CD8^+^ T cells at day 4 post-infection (left), or among total H-2D^b^ GP_33-41_ tetramer^+^ endogenous siIEL CD8^+^ T cells at day 4 post-infection in mice that had not received adoptive transfer of P14 CD8^+^ T cells (right). (D) Expression of Ezh2 among IL-2R*α*^hi^ (blue) and IL-2R*α*^lo^ (gray) P14 siIEL CD8^+^ T cells, as gated in (C). (**E**) Schematic of experimental setup. CD45.1^+^ P14 CD8^+^ T cells were adoptively transferred into CD45.2^+^ congenic hosts one day prior to infection with LCMV. CD45.1^+^ P14 CD8^+^ T cells were collected from the siIEL compartment at day 4 post-infection and sorted based on expression of IL-2R*α* as shown in (C). IL-2R*α*^hi^ and IL-2R*α*^lo^ populations were adoptively transferred into secondary hosts that had been infected with LCMV one day prior to transfer. 7 days later, siIEL and blood P14 CD8^+^ T cells were analyzed by flow cytometry. (**F**) Quantification of the proportion of P14 CD8^+^ T cells present within the siIEL (left) or blood (right) compartments. For siIEL, data were pooled from 2 independent experiments, with mean and SEM of n=4, where each dot represents an individual mouse. For blood, data were pooled from 3 independent experiments, with mean and SEM of n=7 (IL-2R*α*^hi^) or 12 (IL-2R*α*^lo^) hosts, where each dot represents an individual mouse. (**G** and **H**) Quantification of the percent of circulating CD45.1^+^ P14 CD8^+^ T cells derived from IL-2R*α* ^hi^ (blue) or IL-2R*α*^lo^ (gray) day 4 siIEL cells with each indicated phenotype (right), with representative flow cytometry plots shown to the left. Data were pooled from 2 independent experiments, with mean and SEM of n=5 (IL-2R*α*^hi^) or 6 (IL-2R*α*^lo^) hosts, where each dot represents an individual mouse. *P<0.05, **P<0.001 (Student’s two-tailed *t*-test).

In order to determine whether these populations were functionally distinct, P14 CD8^+^CD45.1^+^ T cells were adoptively transferred into CD45.2^+^ hosts infected with LCMV one day later, and the siIEL compartment was harvested at day 4 post-infection for flow cytometric analysis. We found that, in addition to IL-2R*α*^hi^ and IL-2R*α*^lo^ subsets within the early circulating CD8^+^ T cell population, there was also heterogeneity in IL-2R*α* expression among day 4 CD8^+^ T cells within the small intestinal epithelial tissue; notably, this heterogeneity was also evident among endogenous siIEL CD8^+^ H-2D^b^ GP_33-41_ tetramer^+^ cells at day 4 post-infection (Fig. 6C). We also found that protein expression of IL-2R*α* and Ezh2 were correlated (Fig. 6D) and took advantage of this observation to test whether these populations within the tissue have distinct differentiation potentials. CD8^+^IL-2R*α*^hi^ or CD8^+^IL-2R*α*^lo^ cells from the siIEL compartment were FACS-sorted at 4 days post-infection and adoptively transferred intravenously into congenically distinct recipient mice infected with LCMV one day prior to transfer; analysis of blood and siIEL CD8^+^ T cells was performed 7 days later (Fig. 6E). Progeny of day 4 IL-2R*α*^hi^ siIEL CD8^+^ T cells were present at much lower frequencies in both the blood and siIEL compartments of recipient mice than the progeny of their IL-2R*α*^lo^ counterparts, suggesting that the IL-2R*α*^hi^ population might have a lower potential to give rise to both circulating and tissue-resident memory CD8^+^ T cells (Fig. 6F). The inability to detect P14 CD8^+^ T cells within the siIEL compartment of mice that had received IL-2R*α*^hi^ cells precluded further analysis, but cells detected within the blood of these mice expressed much higher levels of KLRG1 and CX3CR1 compared to their counterparts in mice that had received IL-2R*α*^lo^ siIEL CD8^+^ T cells (Fig. 6, G and H). Taken together, these data suggest that, analogous to IL-2R*α*^lo^ and IL-2R*α*^hi^ populations within the circulating CD8^+^ T cell pool early in infection, two functionally distinct CD8^+^ T cell populations are present within the siIEL compartment at day 4 post-infection: a IL-2R*α*^hi^ subset with a more terminally differentiated phenotype that likely represents a transient tissue effector population, and a IL-2R*α*^lo^ subset with higher memory potential that likely contains the precursors of T_RM_ cells. Cluster 20 cells, corresponding to the IL-2R*α*^hi^ population, are more proliferative, exhibit a transcriptional profile that includes factors that are associated with the effector CD8^+^ T cell fate (*Ezh2, Dnmt1*), and are more likely to give rise to circulating terminal effector CD8^+^ T cells. In contrast, Cluster 16 cells, corresponding to the IL-2R*α*^lo^ subset, may be more quiescent and better poised to respond to type I interferons, have greater survival capacity, and have greater potential to give rise to circulating memory CD8^+^ T cells. Taken together, these results demonstrate previously unappreciated transcriptional and functional heterogeneity within the siIEL CD8^+^ T cell pool at multiple states of differentiation.

## Discussion

Recent studies have begun to elucidate key regulators of T_RM_ cell differentiation, function, and survival, such as the transcription factors Blimp1, Hobit, and Runx3. Our single-cell RNA sequencing analyses have identified a number of additional putative regulators of T_RM_ cell differentiation. For example, *Nr4a2*, *Junb,* and *Fosl2* were among the 528 genes that were substantially enriched in siIEL CD8^+^ T cells relative to splenic cells at all time points following infection, and we found that knockdown of *Nr4a2* or *Junb* resulted in impaired T_RM_ cell differentiation. *Junb* and *Fosl2* encode for AP-1 dimerization partners that have been reported to repress T-bet expression in Th17 cells (*46, 53*). Given that downregulation of T-bet expression is important for early establishment of the T_RM_ cell transcriptional program (*17*), it is tempting to speculate that *Junb* and *Fosl2* may regulate T_RM_ cell differentiation by cooperatively regulating T-bet expression. Since *Fosl2* has been previously reported to positively regulate *Smad3* (*53*), a key component of the TGF*β* signaling pathway, *Fosl2* may also promote T_RM_ cell differentiation through effects on TGF*β* signaling.

In addition to identifying specific putative regulators of T_RM_ cell differentiation, our analyses have implicated new pathways that may control aspects of T_RM_ cell biology. For example, we observed that siIEL CD8^+^ T cells exhibited a transcriptional signature indicative of sustained TCR signaling, even after infection had been cleared and antigen was no longer present, suggesting that CD8^+^ T cells within tissues may experience higher basal TCR signals. Since T_RM_ cells have been reported to exhibit a decreased ability to scan for antigens owing to reduced motility relative to circulating CD8^+^ T cells (*54, 55*), such enhanced TCR responsiveness might represent a potential mechanism enabling T_RM_ cells to mount robust protective responses even in the face of limiting amounts of cognate antigen. Moreover, we observed that differentiating siIEL CD8^+^ T cells also exhibited sustained expression of genes encoding inhibitory receptors, including *Ctla4, Tigit*, and *Lag3*, consistent with prior reports (*15, 21*), which may provide a balance against higher basal TCR responsiveness and enable T_RM_ cells to avoid excessive responses that might lead to autoimmune pathology (*56*).

How might T_RM_ cells sustain TCR signaling in the absence of antigen? Since increased plasma membrane cholesterol levels in activated CD8^+^ T cells has been reported to promote TCR clustering and signaling (*57*), it is possible that the increased expression of genes associated with cholesterol synthesis that we observed in siIEL CD8^+^ T cells might contribute to their ability to maintain high levels of TCR signaling. Alternatively, as it has been shown that cytokines can enhance TCR responsiveness (*58*), tonic signals provided by microbiota in the gut microenvironment resulting in cytokine secretion by immune cells might also contribute to the maintenance of genes indicative of sustained TCR signaling in siIEL CD8^+^ T cells in the absence of antigen.

In addition to identifying new pathways regulating T_RM_ cell biology and differentiation, our analyses provide new insights regarding T_RM_ cell ontogeny, which has not been as well studied as that of circulating memory CD8^+^ T cells. It has been previously shown that CD8^+^ T cells can undergo asymmetric division, giving rise to progeny that have distinct tendencies toward the effector or memory fates (*59*). Moreover, single-cell transcriptional profiling analyses (*22, 60*) have demonstrated that CD8^+^ T cells that have undergone their first division in response to microbial infection exhibit divergent transcriptional profiles, with one subpopulation of cells exhibiting a transcriptional burst of genes associated with proliferation and terminal differentiation, while the other subpopulation maintains expression of genes associated with long-lived memory cells. This transcriptional heterogeneity has been linked to functional consequences, as circulating CD8^+^ T cells with high or low expression of IL-2R*α* at the first division or day 3 post-infection exhibited distinct capacities to give rise to long-lived memory cells (*60, 61*), as did CD8^+^ T cells with high or low expression of KLRG1 and IL-7R at the peak of infection (*47, 48*). By contrast, other than the observations that T_RM_ cells are derived from circulating cells lacking high expression of KLRG1 (*15, 18*) and share a common clonal origin as T_CM_ cells (*62*), very little is known about the precursors of T_RM_ cells following their arrival in the tissue.

We found that siIEL CD8^+^ T cells were transcriptionally distinct from splenic CD8^+^ T cells at 4 days post-infection, the earliest time point at which these cells can be detected within the small intestinal tissue (*32*). One interpretation of this finding is that transcriptional changes induced by the local tissue microenvironment occur rapidly upon tissue entry. Alternatively, CD8^+^ T cells that seed tissues may represent pre-committed T_RM_ cell precursors that have already acquired some aspects of the T_RM_ cell transcriptional program. Although there were distinct clusters of splenic CD8^+^ T cells observed at day 3 post-infection, none of these clusters appeared to be transcriptionally similar to siIEL CD8^+^ T cells. This finding suggests that the T_RM_ cell transcriptional program may not be initiated until after the cells have entered the tissue and does not support the hypothesis that pre-committed T_RM_ precursor cells are formed within the circulating CD8^+^ T cell pool. However, it has been shown that priming by DNGR1^+^ dendritic cells in the draining lymph nodes is an important step for T_RM_ cell differentiation in vaccinia virus infection in the skin and influenza A infection in the lung (*63*). These findings suggest that the mechanisms underlying early T_RM_ cell fate specification might differ for localized infections and might even be distinct among different tissues or pathogens. A priming step that specifically induces a T_RM_ cell precursor population that is poised to home to a specific tissue could be particularly advantageous for a localized infection actively occurring within that tissue. In contrast, a more generalized mechanism of T_RM_ cell specification, in which less committed precursor cells enter tissues and differentiate into T_RM_ cells only in response to local tissue signals, might be more permissive for seeding of multiple, diverse tissues, as occurs in a systemic infection. Alternatively, it is possible that both models of T_RM_ cell fate specification could occur simultaneously during an infection, whereby each pathway induces functionally distinct subsets of T_RM_ cells.

In addition to finding substantial transcriptional differences between splenic and siIEL CD8^+^ T cells at day 4 post-infection, we also observed two transcriptionally distinct cell clusters within the siIEL CD8^+^ T cell pool at day 4 post-infection. Cells from one cluster expressed higher levels of genes associated with proliferation and effector differentiation, and exhibited a poor potential to persist in both the circulation and within the tissue when adoptively transferred into new recipients, indicative of a more terminally-differentiated phenotype. Indeed, among the low numbers of transferred cells that were present within the circulation, most exhibited a terminal effector phenotype. By contrast, cells from the other cluster, marked by lower expression of IL-2R*α*, exhibited a greater potential to give rise to both circulating as well as tissue-resident populations upon adoptive transfer. Although it remains unknown whether this heterogeneity is a specific feature of the siIEL compartment or generalizable to T_RM_ cell differentiation in other non-lymphoid tissues, our analyses establish that analogous to the IL-2R*α*^lo^ population present within the circulating CD8^+^ T cell pool early in infection that contains the precursors of circulating memory cells, *bona fide* T_RM_ cell precursors can be found within the IL-2R*α*^lo^ cell subset present within the siIEL compartment at day 4 post-infection. This finding represents an important step in understanding the ontogeny of T_RM_ cells after their arrival in the tissue that will likely inform future studies to identify early determinants of T_RM_ cell fate specification.

Whereas the heterogeneity within the circulating memory CD8^+^ T cell pool is well characterized, heterogeneity among T_RM_ cells is much less clear. Some studies have reported tissue-specific differences in CD69 and CD103 expression by CD8^+^ T cells, but it is unknown whether this heterogeneity represents functionally distinct subsets (*21, 64–66*). Our analyses identify two transcriptionally distinct cell clusters at days 60 and 90 post-infection that exhibit unique functional capacities. One population, distinguished by higher expression of CD28 and IL-7R*α*, expressed high levels of transcripts encoding inflammatory cytokines and exhibited an enhanced ability to produce cytokines in response to restimulation. In addition, this population exhibited higher expression of *Klf2*, which promotes tissue egress and recirculation, and exhibited greater survival potential when adoptively transferred into new recipients. By contrast, the other population, characterized by lower CD28, and IL-7R*α* expression, exhibited a lower potential to give rise to circulating and tissue-resident memory cells when adoptively transferred to new recipients, indicating a poor survival capacity outside the tissue microenvironment. This population also exhibited higher expression of transcripts encoding molecules that promote tissue retention, such as *Cxcr3* and *Klf3* (*67*). Taken together, these findings suggest that one T_RM_ cell subset may exhibit a greater potential to leave the tissue and migrate to a draining lymph node in response to re-infection, where enhanced production of cytokines might promote the activation of circulating memory cells, which can then enter tissues and give rise to secondary T_RM_ cells, as has been recently reported (*68*). By contrast, the second T_RM_ cell subset might be more prone to remain within the tissue and mediate protective responses directly within the tissue microenvironment. Moreover, in addition to revealing functionally distinct subsets within the T_RM_ cell pool, our transcriptional analyses have also elucidated a number of factors regulating the differentiation of these subsets. Analogous to our findings, a recent study provided evidence for distinct subsets of human T_RM_ cells, one with a greater potential for cytokine production and another with a higher proliferative potential in response to TCR stimulation (*69*).

Overall, our work has resulted in a single-cell transcriptomic dataset encompassing the gene-expression patterns of circulating and siIEL CD8^+^ T cells in response to viral infection. Our study reveals a core transcriptional program that is shared between circulating memory and T_RM_ cells in addition to key differences in the kinetics and magnitude of gene expression between these two memory cell subtypes, which should serve as a useful resource for elucidating new genes and pathways regulating T_RM_ cell differentiation. Notably, our analyses demonstrate that CD8^+^ T cells within the siIEL compartment become transcriptionally distinct from circulating cells rapidly upon entry into tissue and identify a subset of early siIEL CD8^+^ T cells enriched for precursors of T_RM_ cells. Moreover, our study reveals previously unappreciated molecular and functional heterogeneity within the T_RM_ cell pool, underscoring the power and necessity of using a single-cell approach. This resource dataset should inform future studies aimed at improving our understanding of CD8^+^ T cell differentiation and function, which may lead to strategies to optimize CD8^+^ T cell responses to protect against microbial infection and cancer.

## Materials and Methods

### Study design

The purpose of this study was to gain a broader understanding of the gene expression patterns that regulate CD8^+^ T cell differentiation and heterogeneity in response to viral infection. To this end, we employed single-cell RNA sequencing on CD8^+^ T cells in the spleen and small intestinal epithelium at various timepoints following viral infection to create a resource dataset analyzing gene expression patterns over time in individual CD8^+^ T cells. Analysis of this dataset revealed a subset of genes whose expression was enriched in CD8^+^ T cells from the tissue, and the role of several of these factors in regulating the differentiation of tissue-resident memory CD8^+^ T cells was validated by flow cytometric analysis of CD8^+^ T cells with shRNA-mediated knockdown of these genes. Additionally, validation of putative subsets of CD8^+^ T cells within the tissue was validated by flow cytometric analysis, including sorting of individual putative subsets and assessment of function following *ex vivo* restimulation or adoptive transfer into secondary hosts. Information regarding the sample size and number of replicates for each experiment can be found in the relevant figure legend. All findings were successfully reproduced, except for measurements of CD28 expression in P14 intraepithelial T cells with knockdown of *Pnrc1* and analyses of *Ddx5*^-/-^ CD8^+^ T cells, which were performed once. No sample size calculations were performed; sample sizes were selected based on previous studies performed in our lab. Mice were randomly allocated into groups prior to adoptive transfer of sorted IL-2R*α*^hi^ or IL-2R*α*^lo^ intraepithelial P14 T cells (Fig. 6). For single-cell RNA sequencing experiments, mice were randomly selected for P14 T cell harvesting at specific time points post-infection. Randomization and blinding are not relevant to shRNA knockdown experiments, or experiments involving co-transfer of *Ddx5*^-/-^ and WT CD8^+^ T cells, since P14 T cells transduced with retrovirus encoding shRNA against genes of interest and congenically distinct P14 T cells transduced with control retrovirus, or *Ddx5*^-/-^ and WT CD8^+^ T cells, were co-transferred into the same animals (Fig. 2 and Fig. 5). For assessment of phenotype of adoptively transferred sorted IL-2R*α*^hi^ or IL-2R*α*^lo^ intraepithelial P14 T cells (Fig. 6), the investigator was aware of the cell type transferred into each recipient. No data were excluded from analysis, except for recipient mice that had rejected adoptively transferred P14 CD8^+^ T cells.

### Mice

All mice were housed under specific pathogen-free conditions in an American Association of Laboratory Animal Care-approved facility at the University of California, San Diego (UCSD), and all procedures were approved by the UCSD Institutional Animal Care and Use Committee. Wild-type C57BL6/J (CD45.2^+^) and P14 TCR transgenic (CD45.1 or CD45.1.2^+^, both maintained on a C57BL6/J background) mice were bred at UCSD or purchased from Jackson Laboratories. *Ddx5^fl/fl^* mice were obtained from Dr. Frances Fuller-Pace’s laboratory (University of Dundee) and have been previously described (*70*). To obtain congenically distinct P14 *Ddx5^fl/fl^*CD4-Cre^+^ and P14 *Ddx^fl/fl^*CD4-Cre^-^ mice, *Ddx5^fl/fl^* mice were crossed to P14 CD4-Cre^+^ mice (either CD45.1 or CD45.1.2^+^). All mice were used from 6-9 weeks of age, male mice were used as recipients, and male or female mice were used as donors in adoptive transfer experiments.

### Antibodies, flow cytometry, and cell sorting

Cells were stained for 10 minutes on ice with the following antibodies: V*α*2 (B20.1), CD8*α* (53-6.7), CD8*β* (YTS156.7.7), CD45.1 (A20), CD45.2 (104), CD44 (1M7), CX3CR1 (SA011F11), CD127 (A7R34), CD27 (LG.3A10), CD69 (H1.2F3), CD103 (2E7), CD28 (37.51), CD25 (PC61), and KLRG1 (2F1/KLRG1) all purchased from Biolegend. In some experiments, cells were stained with H-2D^b^ GP_33-41_ tetramer (obtained from the NIH Tetramer Core) for 1 hour at room temperature prior to staining with cell surface antibodies. Samples were then stained in Fixable Viability Dye eFluor780 (Thermo Fisher Scientific) or Ghost Violet 510 (Tonbo Biosciences) at 1:1000 on ice for 10 minutes. For experiments with retroviral transduction of P14 T cells, cells were then fixed in 2% paraformaldehyde (Electron Microscopy Services) on ice for 45 minutes. For staining for Ezh2, cells were fixed and permeabilized using the FoxP3/Transcription Factor Staining Buffer Kit (Thermo Fisher) following staining with viability dye, prior to incubation with anti-Ezh2 antibody (11/Ezh2, BD Pharmingen) for 8 hours at 4°C. For assessment of cytokine production, cells were cultured in the presence of LCMV GP_33-41_ peptide (GenScript) and Protein Transport Inhibitor Cocktail (ThermoFisher Scientific) for 3 hours at 37°C. Following cell surface and viability staining, cells were fixed and permeabilized using BD Cytofix/Cytoperm (BD Biosciences) for 30 minutes at room temperature prior to staining with anti-IFN*γ* (XMG1.2), TNF*α* (MP6-XT22), and IL-2 (JES6-5H4) antibodies (all from Biolegend) for 30 minutes on ice. For analysis, all samples were run on an Accuri C6, LSRFortessa, or LSRFortessa X-20 (BD Biosciences). For sorting, all samples were run on an Influx, FACSAria Fusion, or FACSAria2 (BD Biosciences). BD FACS DIVA software (BD Biosciences) was used for data collection, and Flowjo software (Treestar) was used for analysis of flow cytometry data.

### Naïve T cell transfer and infection

Splenocytes were collected from naïve CD45.1^+^ or CD45.1.2^+^ P14 mice and stained with antibodies against V*α*2, CD8, and CD45.1. 1×10^5^ V*α*2^+^CD8^+^CD45.1^+^ cells were adoptively transferred into congenically distinct wild-type recipients one day prior to infection with 2×10^5^ plaque-forming units (PFU) of LCMV-Armstrong, injected intraperitoneally. For experiments where IEL were collected at 4 days post-infection, 5×10^5^ V*α*2^+^CD8^+^CD45.1^+^ P14 T cells were transferred.

### CD8^+^ T cell isolation

For isolation of CD8^+^ T cells from spleen, spleens were collected and dissociated to yield a cell suspension prior to treatment with Red Blood Cell Lysing Buffer Hybri-Max (Sigma). For isolation of CD8^+^ T cells from the small intestinal epithelium, Peyer’s patches were removed and the tissue was cut longitudinally and washed of luminal contents. The tissue was then cut into 1cm pieces that were incubated while shaking in DTE buffer (1ug/ml dithioerythritol (Thermo Fisher Scientific) in 10% HBSS and 10% HEPES bicarbonate) at 37°C for 30 minutes. Cells within the supernatant were collected and passed through a 44/67% Percoll density gradient to enrich for lymphocytes.

### 10X Genomics library preparation and sequencing

Activated P14 T cells (CD8^+^V*α*2^+^CD45.1^+^CD44^+^) were sorted from the spleen or siIEL and resuspended in PBS+0.04% (w/v) bovine serum albumin. Approximately 10,000 cells per sample were loaded into Single Cell A chips (10X Genomics) and partitioned into Gel Bead In-Emulsions (GEMs) in a Chromium Controller (10X Genomics). Single cell RNA libraries were prepared according to the 10x Genomics Chromium Single Cell 3’ Reagent Kits v2 User Guide, and sequenced on a HiSeq4000 (Illumina).

### Generation of shRNA-encoding retrovirus, transduction of CD8^+^ T cells, and adoptive transfer for analysis of gene knockdown

shERWOOD-designed UltramiR sequences targeting *Junb, Nr4a2*, *Pnrc1,* or *Zfp36l2* (knockdown, KD) or control (non-targeting) constructs in LMP-d Ametrine vector were purchased from transOMIC technologies. To generate retroviral particles, 293T HEK cells were plated in 10cm plates one day prior to transfection and individually transfected with 10ug of each shRNA retroviral construct and 5ug pCL-Eco using TransIT-LT1 (Mirus). Retroviral supernatant was collected and pooled at 48 and 72 hours post-transfection, and *in vitro* activated P14 T cells were transduced with retrovirus encoding KD shRNA, and congenically distinct *in vitro* activated P14 T cells were transduced with control retrovirus. To activate P14 T cells *in vitro*, lymphocytes were collected from spleens and lymph nodes of naïve CD45.1^+^ and CD45.1.2^+^ P14 TCR transgenic mice and negatively enriched for CD8^+^ T cells using the CD8a^+^ T Cell Isolation Kit and LS MACS Columns (Miltenyi Biotec). 1×10^6^ CD8^+^ T cells were plated per well in 48 well flat-bottom plates, pre-coated with 100ug/ml goat anti-hamster IgG (H+L, Thermo Fisher Scientific) followed by 5ug/ml each anti-CD3 (clone 3C11, BioXCell) and anti-CD28 (clone 37.51, BioXCell). Eighteen hours after activation, cells were transduced with retroviral supernatant supplemented with 8ug/ml polybrene (Millipore) by spinfection for 90 minutes at 900rcf at room temperature. Following spinfection, retroviral supernatant was removed and replaced with culture medium (Iscove’s modified Dulbecco’s medium+10% fetal bovine serum (v/v)+2mM glutamine+100U/ml penicillin+100ug/ml streptomycin+55mM *β*-mercaptoethanol) and cells were rested for 2 hours at 37°C. Cells transduced with each individual construct were then pooled, washed three times with PBS, counted, and mixed to obtain a 1:1 ratio of transduced KD and transduced control cells, based on previously tested transduction efficiency of each retrovirus. 5×10^5^ total P14 T cells were then adoptively transferred into congenically distinct hosts that were infected with 2×10^5^ PFU LCMV-Arm one hour later. 1×10^6^ cells from this mixture were returned to culture with recombinant IL-2 (100U/ml), and analyzed 18 hours later by flow cytometry to determine the input ratio of transduced KD:control P14 T cells. 22-26 days post-infection, splenocytes and siIEL were collected as described above, and analyzed by flow cytometry to determine the ratio of KD:control cells within each subset of transduced P14 T cells.

### Statistical analysis of flow cytometry data

Statistical analysis of flow cytometry data was performed using Prism software (Graphpad). P values of <0.05 were considered significant. Statistical details for each experiment can be found within the relevant figure legends.

### Quantitative and statistical analysis of scRNA-seq data

#### Single-cell RNA-seq mapping

Reads from single-cell RNA-seq were aligned to mm10 and collapsed into unique molecular identifier (UMI) counts using the 10X Genomics Cell Ranger software (version 2.1.0). All samples had sufficient numbers of genes detected (>1000), a high percentage of reads mapped to the genome (>70%), and sufficient number of cells detected (>1000).

### Cell and gene filtering

Raw cell-reads were then loaded to R using the cellrangerRkit package. The scRNA-seq dataset was then further filtered based on gene numbers and mitochondria gene counts to total counts ratio. To ensure that memory requirements for all downstream analyses did not exceed 16Gb and that the samples with more cells would not dominate the downstream analysis, we randomly selected a portion of the cells that passed filtering for downstream analysis. We randomly selected ∼2000 cells from each library for downstream analysis. After cell filtering and sampling, we filtered genes by removing genes that did not express > 1 UMI in more than 1% of the total cells.

### Single-cell RNA-seq dataset normalization and pre-processing

Five cell-gene matrices were generated:

1. Raw UMI matrix.
2. UPM matrix. The raw UMI matrix was normalized to get UMIs per million reads (UPM), and was then log2 transformed. All downstream differential analysis was based on the UPM matrix. The prediction models were also based on the UPM matrix, as other normalizations are very time-consuming for large datasets.
3. MAGIC matrix. UPM matrix was further permuted by MAGIC (van Dijk et al., 2018) R package Rmagic 1.0.0 was used, and all options were kept as default. MAGIC aims to correct the drop-out effect of single-cell RNA-seq data; thus, we used MAGIC-corrected matrix for visualizing the gene expression pattern rather than using the UPM matrix. All gene expression heatmaps and gene expression overlaid on TSNE plots were based on the MAGIC matrix.
4. Super cell matrix. We merged 50 cells to create a ‘super’ cell and used the super cell matrix as the input for WGCNA and cell type annotation analysis. This approach enabled us to bypass the issue of gene dropouts with scRNA-seq and is equivalent to performing WGCNA on thousands of pseudo-bulk samples. We first calculated the mutual nearest neighbor network with k set to 15, and then cells that were not mutual nearest neighbors with any other cells were removed as outliers. We randomly selected ‘n’ cells in the UPM matrix as the seed for super cells. The expression of each super cell was equal to the average expression of its seed and the 50 nearest neighbor cells of its seed. We derived 7400 super cells from the dataset, so each single cell was covered ∼10 times.

### Single-cell RNA-seq dataset dimension reduction

Top variable genes, PCA, and tSNE were calculated by Seurat version 2.3.4 functions: FindVariableGenes, RunPCA, and RunTSNE (Butler et al., 2018). Only the top 3000 genes were considered in the PCA calculation and only the top 25 principal components (PCs) were utilized in tSNE. Louvain clustering was performed by Seurat’s FindClusters function based on the top 25 PCs, with resolution set to 2.

### Differential gene expression analysis

Differentially expressed (DE) genes were identified by performing pairwise comparison using two-sided Wilcox test and the FindAllMarkers function of Seurat. The threshold for DE genes was p-value < 0.05 and an absolute fold-change > 2.

### WGCNA analysis

We performed WGCNA^6^ (version 1.63) analysis on spleen cells alone, siIEL cells alone, or all cells considered together. The super cell matrices were used as the input to boost performance. Only the top 5000 variable genes were considered in this analysis. SoftPower was set to 9 and the signed adjacency matrix was calculated for gene module identification. Genetree clustering and eigengene clustering were based on average hierarchical clustering. Module cut height was set to 0.1. Gene ontology (GO) analysis of the gene module was performed by compareClusterfunction from R package clusterProfiler 3.2.14. Reference database was org,Hs.eg.db 3.4.0 from Bioconductor and options were fun = “enrichGO”, pAdjustMethod = “fdr”, pvalueCutoff = 0.01 and qvalueCutoff = 0.05.

### Annotating single-cells with bulk RNA-seq signatures

The log2 TPM data from bulk RNA-seq datasets were compared with the scRNA-seq super cell matrix. Bulk cell population RNA-seq samples were first grouped into different sets according to their mutual similarities. For each bulk RNA-seq sample set, the mean expression was first calculated. The 1^st^ correlation was calculated between all the super cells and the mean expression from the bulk RNA-seq dataset. Based on the distribution of the 1^st^ correlation, we were able to identify a group of super cells that were most similar to the mean expression of the bulk sample. To further identify the small differences between bulk RNA-seq expression within a given set, we removed the set mean from the bulk RNA-seq and the mean from the most similar group of super cells, and then calculated the 2^nd^ correlation between the super cells and bulk RNA-seq. Based on the 2^nd^ correlation, we annotated the super cells with each bulk sample label.

## Acknowledgements

We thank members of the Chang, Yeo, and Goldrath laboratories for technical advice, helpful discussion, and critical reading of the manuscript.

## Author contributions

N.S.K, J.J.M., K.D.O, M.S.T., C.E.W., B.S.B., C.M., A.W.G., G.W.Y., and J.T.C. designed experiments and analyzed data; Z.H. and G.W.Y. performed computational analyses; N.S.K, J.J.M., K.D.O, M.S.T., C.E.W., T.L.L., J.N.K., J.G.O., T.T., and L.K.Q. performed experiments; W.J.H. provided key materials; A.W.G., G.W.Y., and J.T.C. supervised the study; and N.S.K. and J.T.C. wrote the first draft of the manuscript. All authors reviewed and edited the manuscript

## Data and materials availability

The single-cell RNA sequencing data presented are available for download on GEO (data repository accession number is pending). Requests for data or resources presented in this study should be directed to the corresponding author.

## Supplementary Materials

**Fig. S1.**
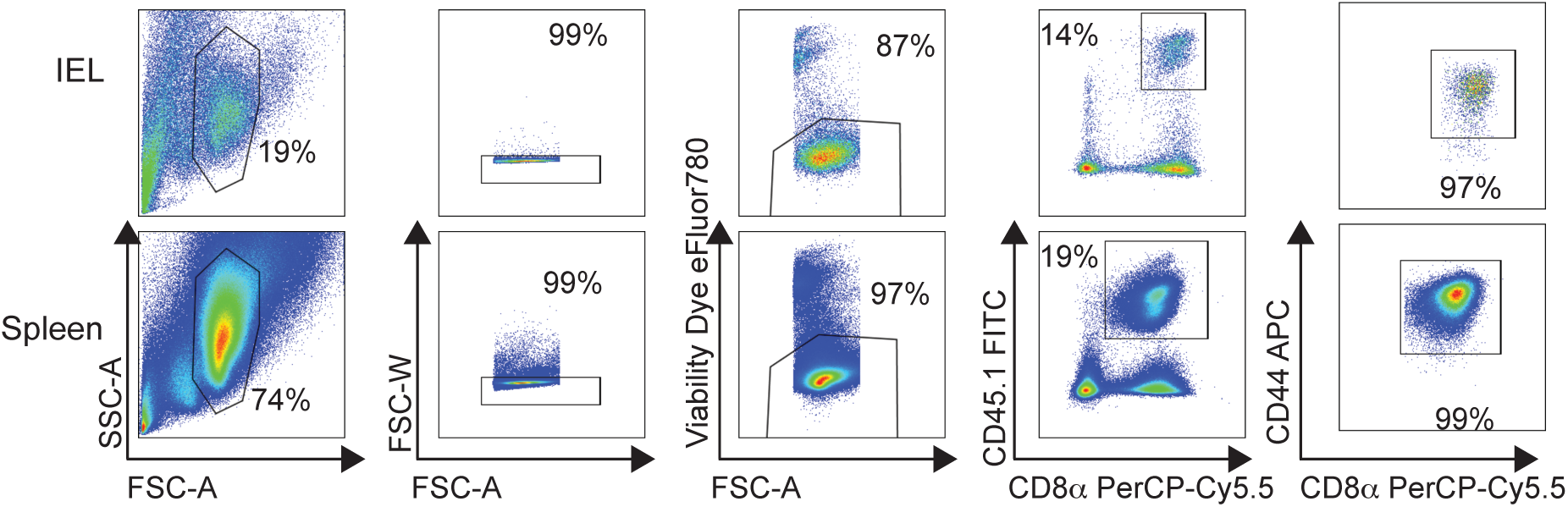
Sorting of P14 CD8^+^ T cells from spleen and siIEL for scRNA-seq. CD45.1^+^ P14 CD8^+^ T cells were adoptively transferred into CD45.2^+^ congenic hosts one day prior to infection with LCMV-Armstrong. Spleens and the siIEL compartments were harvested at the indicated time points, and P14 CD8^+^ T cells were isolated by FACS sorting according to the gating strategy shown for the siIEL compartment (top row) and spleen (bottom row).

**Fig. S2.**
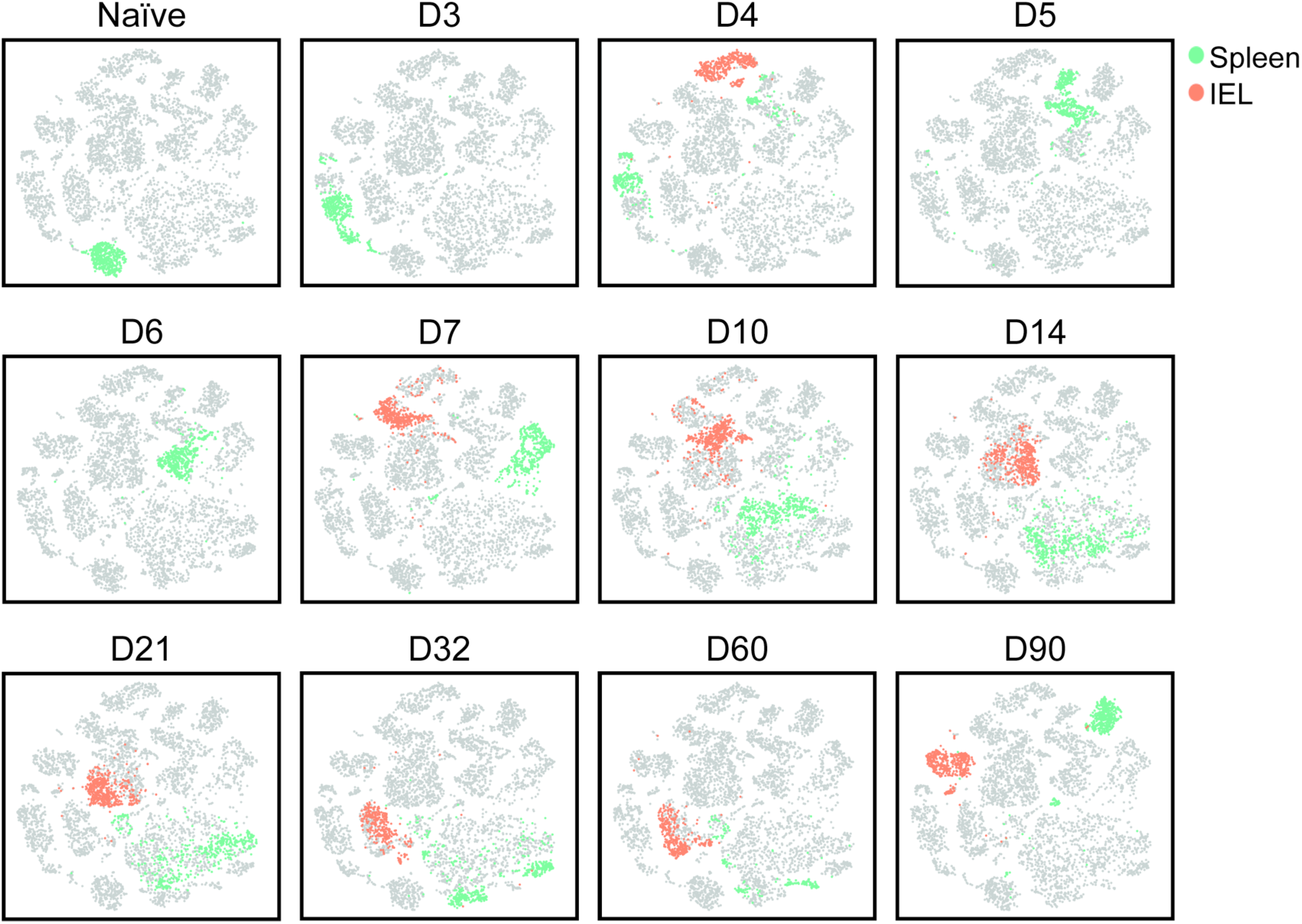
Circulating and siIEL CD8^+^ T cells are transcriptionally distinct at all time points post-infection. Unsupervised t-distributed stochastic neighborhood embedding (tSNE) analyses. Each panel represents an individual time point; teal color represents splenic CD8^+^ cells at the indicated time point, coral color represents siIEL CD8^+^ T cells at the indicated time point, and gray color represents CD8^+^ T cells from other time points.

**Fig. S3.**
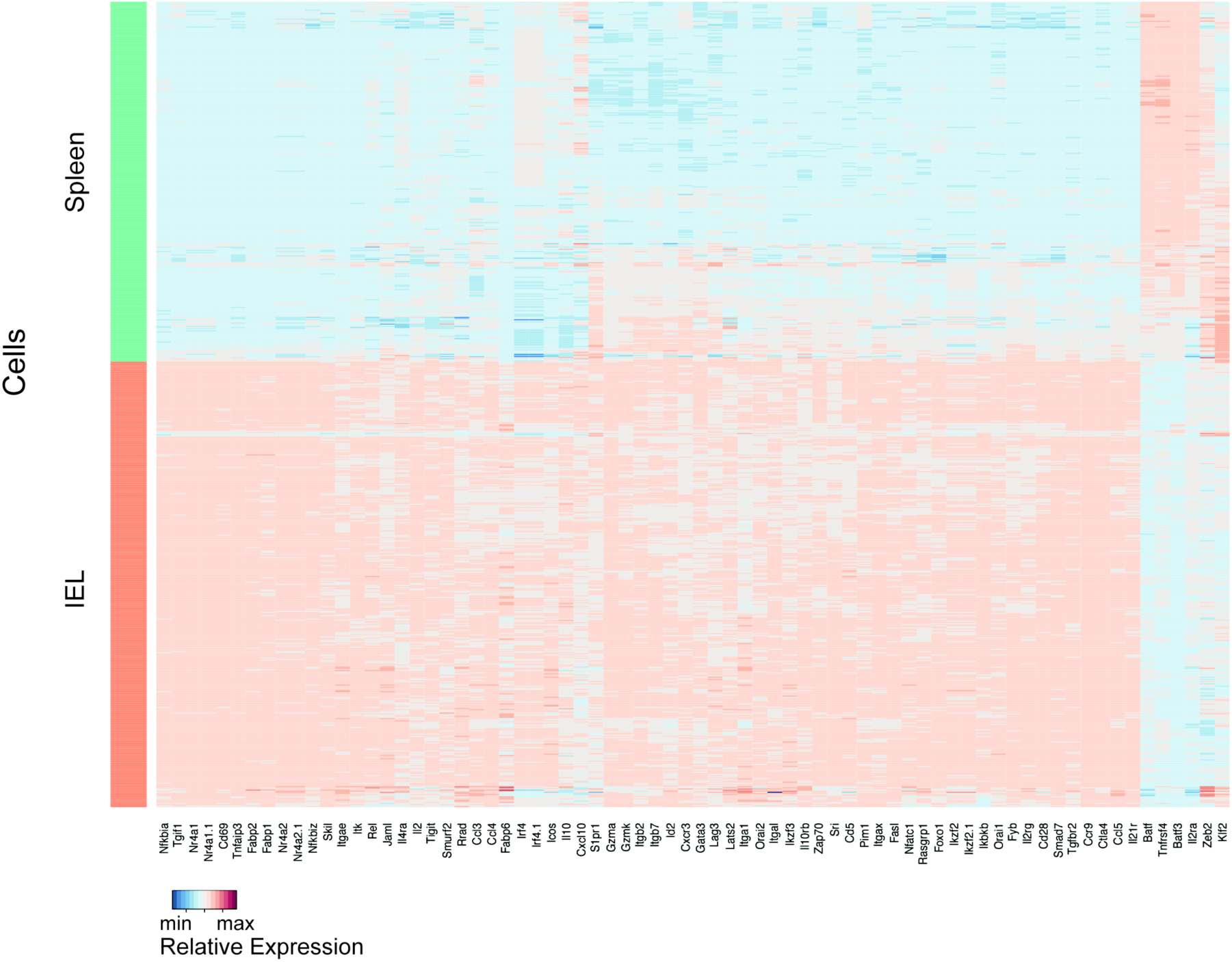
Differential gene expression of circulating and siIEL CD8^+^ T cells at day 4 post-infection. Differential gene expression of selected genes in day 4 splenic (teal) and siIEL (coral) CD8^+^ T cells, represented as expression relative to the mean expression among all cells, where each row represents an individual cell and each column represents an individual gene.

**Fig. S4.**
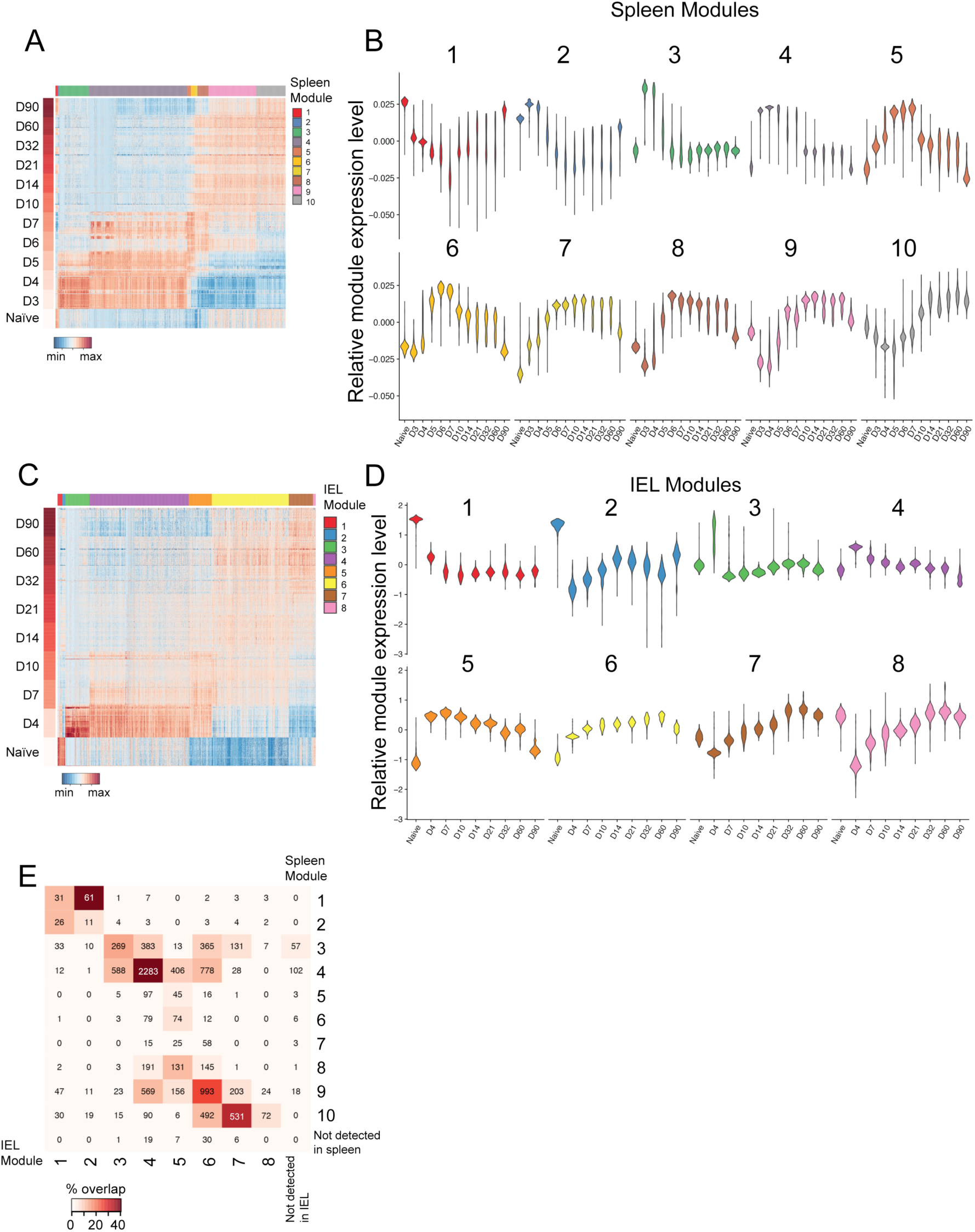
Shared core transcriptional program among splenic and siIEL CD8^+^ T cells. (**A** and **C**) Gene-expression of single CD8^+^ T cells (as in Figure 1A) within the spleen (A) or siIEL compartment (C) responding to infection over time, where each row represents an individual cell, grouped by time point, and each column represents an individual gene, grouped by module, represented as expression relative to the mean expression among all cells. Weighted gene co-expression network analyses of splenic and siIEL CD8^+^ T cells considered separately were performed to derive gene modules. (**B** and **D**) Violin plots depicting the representation of each gene module among single CD8^+^ T cells over time in the spleen (C) or siIEL compartment (D), relative to the mean representation among all cells within that tissue. (**E**) Overlap of genes within distinct Spleen and IEL Modules, where the number in each square represents the number of genes shared between the two modules and the color represents the proportion of the total genes in the two modules that are shared.

**Fig. S5.**
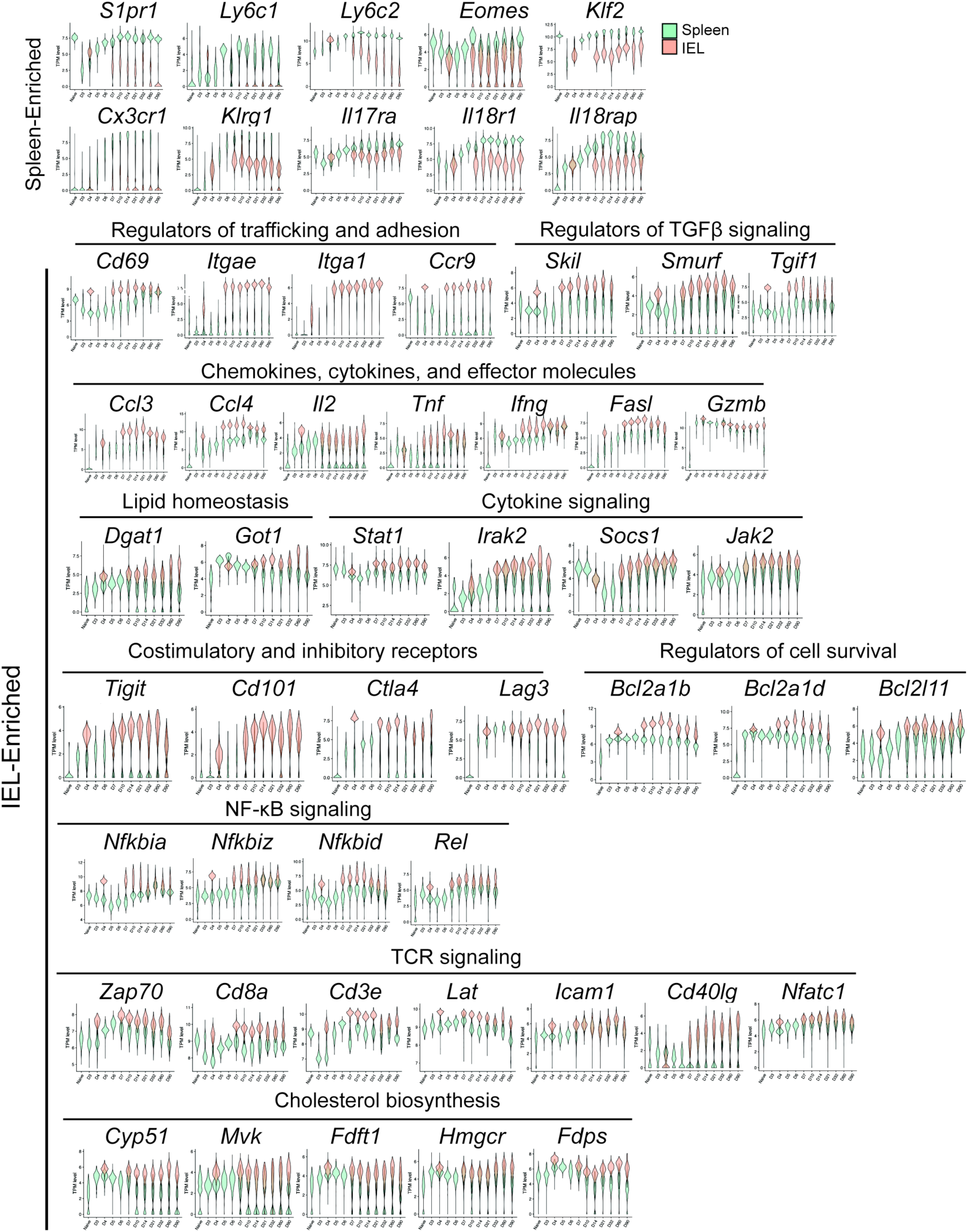
Components of the T_RM_ cell-enriched transcriptional signature. Violin plots depicting expression patterns of known or putative regulators of T_RM_ cell differentiation (represented as transcripts per million, TPM), selected from spleen-enriched or siIEL-enriched modules, among single splenic (teal) or siIEL (coral) CD8^+^ T cells over time.

**Fig. S6.**
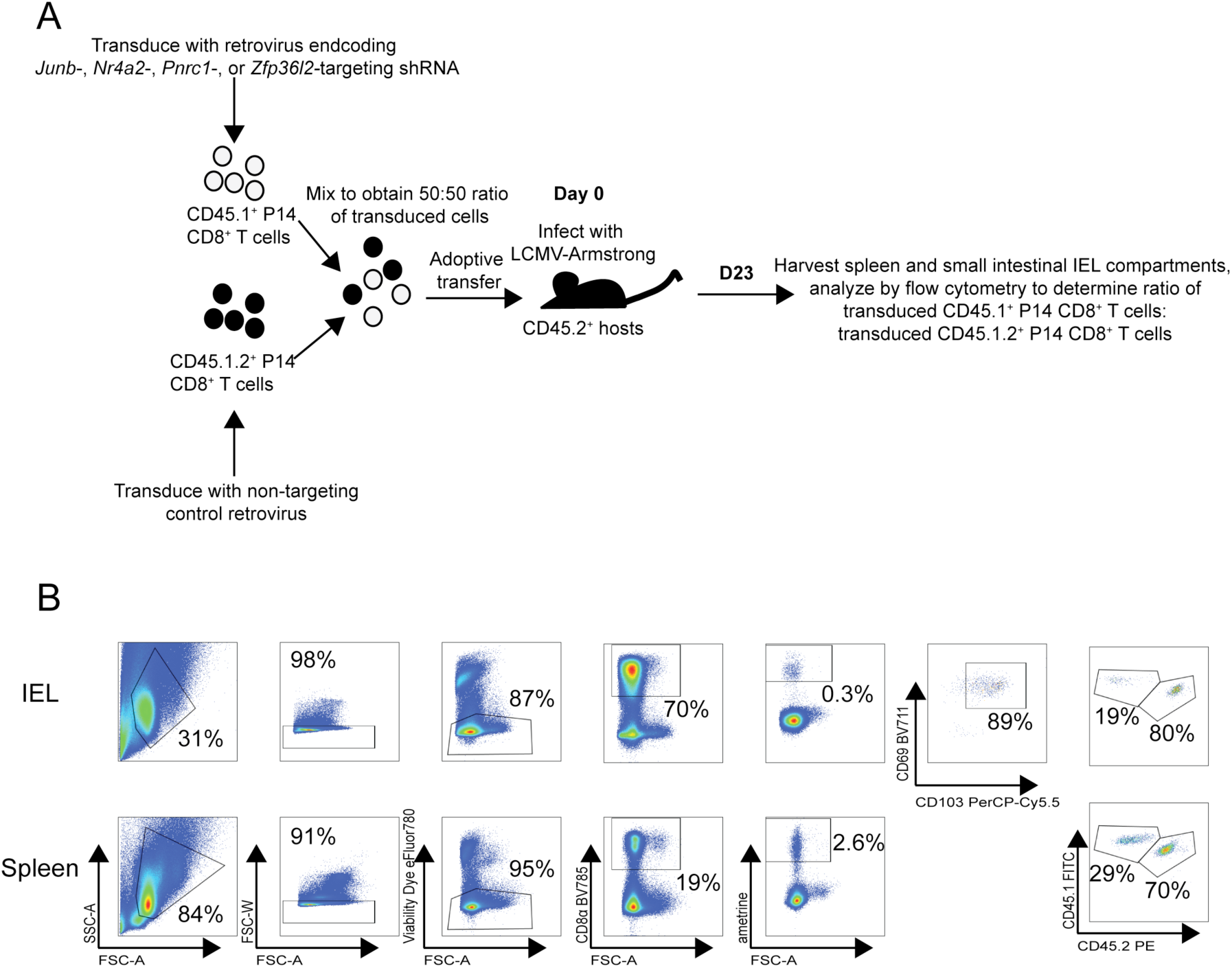
shRNA knockdown of putative regulators of T_RM_ cell differentiation. (**A**) Schematic of experimental setup. CD45.1^+^ P14 CD8^+^ T cells were transduced with retrovirus encoding shRNAs targeted to *Nr4a2* or *Junb* (knockdown, KD), and mixed with CD45.1.2^+^ P14 T cells transduced with shRNA encoding control (non-target) shRNA at a 1:1 ratio of KD:non-target cells prior to adoptive transfer into CD45.2^+^ hosts that were subsequently infected with LCMV. 22-26 days later, splenic and siIEL CD8^+^ T cells were analyzed by flow cytometry. (**B**) Gating strategy for assessing impact of knockdown on tissue-resident (top row) or circulating (bottom row) CD8^+^ T cell memory populations.

